# Cold hardiness dynamics predict budbreak and associated low temperature threats in grapevine

**DOI:** 10.1101/2025.01.09.632158

**Authors:** Francisco Campos-Arguedas, Erica Kirchhof, Michael G. North, Jason P. Londo, Terry Bates, Cornelis van Leeuwen, Agnès Destrac-Irvine, Benjamin Bois, Al P. Kovaleski

**Author notes:** Corresponding author: Al P. Kovaleski. This work represents equal efforts by these authors (Francisco Campos-Arguedas, Erica Kirchhof). **Author contributions:** APK and JPL conceived this work. APK, FC-A, EK, and MGN developed the methodology. All authors contributed to data collection and curation. FC-A, EK, APK and MGN analyzed data and produced figures. FC-A and EK drafted the original manuscript under the supervision of APK, with reviews and contributions from MGN, JPL, TB, CVL, ADI and BB. FC-A and EK contributed equally to this work.

## Abstract

Temperate woody perennial plants form buds that develop into leaves or flowers that emerge in the following growth season. To survive winter, dormant buds must attain cold hardiness, and timely lose it in spring to break bud while avoiding damage from low temperatures and late frosts. Here, we use a process-based model to predict bud cold hardiness of three grapevine varieties (*V. vinifera* ‘Cabernet-Sauvignon’ and ‘Riesling’, and *V. hybrid* ‘Concord’) from historical temperature records of eight different locations in North America and Europe (n=329). Based on those predictions, and thresholds of cold hardiness at budbreak from literature, timing of budbreak was extracted. Despite the model being untrained to budbreak data, the cold hardiness model resulted in good predictions (RMSE=7.3d) that were further improved based on expected delays from cold damage (RMSE=7.2d). Both increasing and decreasing trends in freeze damage risk were predicted with increasing temperature, depending on the range of mean dormant season temperature (MDST; 1 Nov–30 Apr) in each location. Predictions in timing of spring phenology in relation to MDST also showed warming to advance (MDST<10°C) or delay (MDST>10°C) budbreak. Cold hardiness dynamics represents an advancement in phenological modeling that provides information for the entirety of the dormant season, as well as budbreak.

## Introduction

Fluctuations and extremes in temperature, photoperiod, precipitation, and numerous other environmental variables determine the distribution and adaptation of species worldwide (Inouye, 2022). In temperate ecosystems, low temperature is considered one of the most significant factors influencing the distribution of woody perennial species, particularly near the cold edge of their distribution range (Chamberlain & Wolkovich, 2021; Körner, 2021; Qian *et al*., 2022). Seasonal fluctuations in temperature drive phenological events – the timing of which impacts the length of the growing season, overall productivity, and nutrient cycles (Richardson *et al*., 2013; Gallinat *et al*., 2021; Marquis & Lajoie, 2024). Understanding the timing of life cycle events such as budbreak has significant practical and ecological implications, as it informs management practices and climate adaptation strategies (Richardson *et al*., 2013). Predicting timing of spring phenology is also an essential component of Earth system models, which help us understand and forecast ecosystem responses to climate change, including carbon sequestration potential of plants (Cleland *et al*., 2012). Despite the importance of budbreak, accurately forecasting this critical phase remains a challenge due to our general lack of understanding of temperature responses associated with this phase in plants (Ding *et al*., 2024). This is particularly relevant in the face of rapidly changing climates and the emergence of non-analogous environmental conditions (Darbyshire et al., 2013; Piao et al., 2019; Wheeler et al., 2024).Therefore, new models that deviate from standard and incorporate physiologically derived temperature responses may improve our predictive ability for budbreak.

Buds are formed in the late summer and early fall and remain dormant [with little developmental activity (Goeckeritz *et al*., 2023)] throughout winter prior to growth resumption in the spring. Typical phenological models use the experimentally demonstrated negative relationship between accumulation of chill (‘chilling’) – cumulative exposure to low temperatures – and the accumulation of heat (‘forcing’) – cumulative exposure to warm temperatures – to predict budbreak (Wang *et al*., 2020). However, there is likely an overlap in temperature ranges in which chilling and forcing occur (Flynn & Wolkovich, 2018). Therefore, the timing of transition from one phase to another can influence whether mid-dormant season temperatures are contributing to one process or the other (or occasionally both). In most phenology models, the transition from dormant to a growth-responsive state is either classified as sequential or overlapping (Chuine, 2000). In a sequential approach, both chill and heat accumulation occur in separate phases – chilling and forcing – without accounting for their interaction (Ashcroft *et al*., 1977). During the first phase, dormancy release is modulated by exposure to low temperatures (chilling accumulation) (Egea *et al*., 2003). The second phase involves exposure to warm temperatures – accumulation of growing degree days/hours (Fernandez *et al*., 2021). However, delays in the transition between the chilling and heat-accumulation phases can significantly influence the contribution of chill and heat units, and in many cases the timing of this transition is arbitrarily defined or imposed by the model (Egea et al., 2003; Harrington et al., 2010). These models are therefore inherently vulnerable to shifting climatic conditions. Warming temperatures may increase the time exposed to forcing conditions in winter and spring, which would by itself lead to advances in spring phenology. However, the warming conditions will also decrease chill accumulation in fall and spring, which can delay spring phenology. Therefore, variable outcomes in phenological timing can be expected based on future warming (Chamberlain *et al*., 2019; Fernandez *et al*., 2023). As a result, there is considerable interest in developing generalizable models which can accurately describe phenology under a wide range of conditions, especially considering the growing body of evidence supporting both phenological advancement and delay in woody perennials due to climate warming (Hassan *et al*., 2024) and urbanization (Li *et al*., 2019).

Previous work has shown that budbreak can be significantly influenced by cold hardiness dynamics (Kovaleski *et al*., 2018; Kovaleski, 2022; North *et al*., 2022; North & Kovaleski, 2024). Cold hardiness is a plastic trait within a genotype mainly influenced by air temperature, with gains (acclimation) occurring in autumn and early winter and losses (deacclimation) in late winter and early spring. Rates of acclimation, deacclimation, and maximum levels of cold hardiness achieved in a given location vary by genotype and yearly weather conditions (Ferguson *et al*., 2011, 2014; Kovaleski *et al*., 2018, 2023; North *et al*., 2021). Thus, cold hardiness has been shown to be an important barrier to plant survival in cold regions, contributing to the suitability of different species and ecotypes to different climates (Körner *et al*., 2016; Meier *et al*., 2018; De Rosa *et al*., 2021; Körner, 2021; Baranger *et al*., 2024). Deacclimation always precedes budbreak in springtime, with most cold hardiness lost by budbreak (Lenz *et al*., 2013; Vitra *et al*., 2017; Chamberlain & Wolkovich, 2021; Hillmann *et al*., 2021; Kovaleski, 2022; North *et al*., 2022). Effects of cold damage have already been linked to delays in spring phenology within the same year (Qiu *et al*., 2024) and the subsequent year (Wang *et al*., 2025), Altogether, this indicates that measuring or predicting the cold hardiness phenotype and its dynamics may be useful for delivering predictions of budbreak for species across varying climates.

The focus of this study was to test a process-based cold hardiness model for grapevine, NYUS.1 (Kovaleski *et al*., 2023), for predicting budbreak phenology and exploring other climate-related growth risks for three different cultivars across eight locations in Europe and North America. Grapevine (*Vitis* spp.) is both a horticulturally and historically significant crop grown throughout the world, with areas of cultivation present on all continents except Antarctica. Critically, viticultural regions also contain the same clonally propagated genotypes (i.e., cultivars or varieties), facilitating evaluation of plasticity across different climates. The considerable genotypic variability in climatic adaptation among commonly grown grapevines (Wolkovich *et al*., 2018) makes it an interesting model to understand the connections between cold hardiness predictions and estimation of budbreak timing. Without reparameterization, the NYUS.1 model was applied to historical daily temperature records in each location and predictions were validated using longstanding records of phenology observations. Additionally, we used cold hardiness trajectories to predict instances of freeze damage across cultivars and locations, validating these predictions based on newspaper and extension records for one location. Using model outputs, we assessed patterns in freeze risk over both thermal and temporal scales, and estimated phenological sensitivity as a product of dormant season temperatures.

## Materials and Methods

### Phenological Records

The phenology dataset used in this study includes three grapevine cultivars: *Vitis vinifera* cvs. Cabernet-Sauvignon and Riesling, and *Vitis labruscana* cv. Concord (hereon referred to by cultivar name only). These cultivars were used as the only three for which parameters are available for the NYUS.1 grapevine cold hardiness model (Kovaleski et al., 2023). These cultivars have relatively different cold hardiness dynamics patterns, and are representative of different groups of cultivars: Cabernet-Sauvignon is a warm-climate variety, with low cold hardiness and low deacclimation rates; Riesling is a cool climate variety, with intermediate cold hardiness (or high within *V. vinifera*), and moderate deacclimation rates; Concord is a Northern hybrid, with high cold hardiness and high deacclimation rates (Londo & Kovaleski, 2025). Phenology data was retrieved from eight different locations across Europe and North America (**Fig. 1**): Niagara-on-the-Lake, ON, CA (Hébert-Haché *et al*., 2021), Portland, NY, US, Lewisburg, PA, US (Persico *et al*., 2021), Bordeaux, FR (Maury *et al*., 2023), Bergheim, FR, Montreuil-Bellay, FR (Duchêne, 2019), Marseillan-Plage, FR (Maury *et al*., 2023), and Veitshöchheim, DE (Bock *et al*., 2011). For Bordeaux, FR, we included additional phenology data collected in the same plot for the years 2012-2023. For Veitshöchheim, DE, budbreak data were extracted from Bock et al. (2011) Figure 2a in RStudio using metaDigitiseR version 1.0.1 (Pick *et al*., 2019).

**Figure 1.**
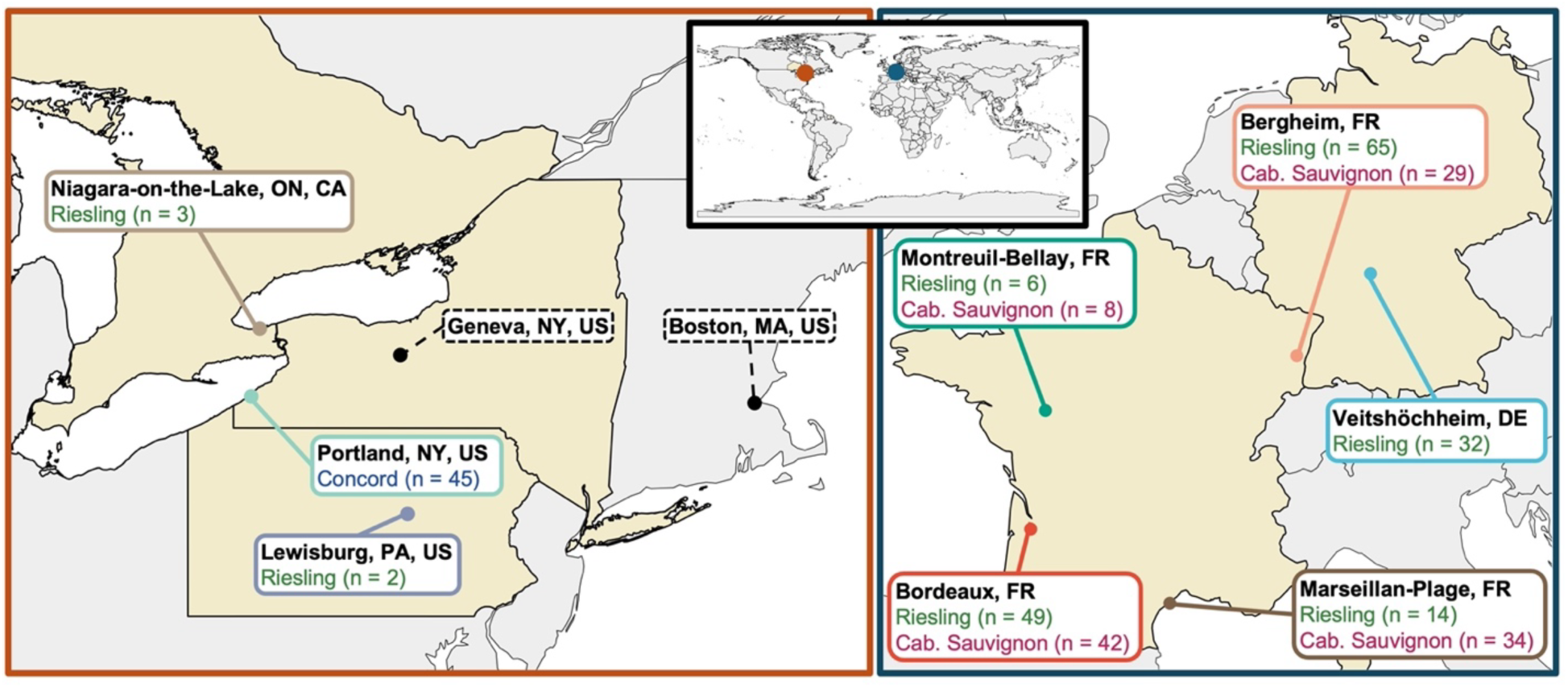
Locations and cultivars in the phenology dataset and cold hardiness model development. The phenology dataset used here contains eight locations: four in France (Bergheim, Bordeaux, Marseillan-Plage, and Montreuil-Bellay), one in Germany (Veitshöchheim), two in the United States (Lewisburg, PA, and Portland, NY), and one in Canada (Niagara-on-the-Lake, ON). The NYUS.1 cold hardiness model was developed using grapevine bud cold hardiness data from Geneva, NY (Kovaleski et al., 2023), and used dormancy parameters from 15 unrelated woody perennial species grown in Boston, MA – both in the United States.

**Figure 2.**
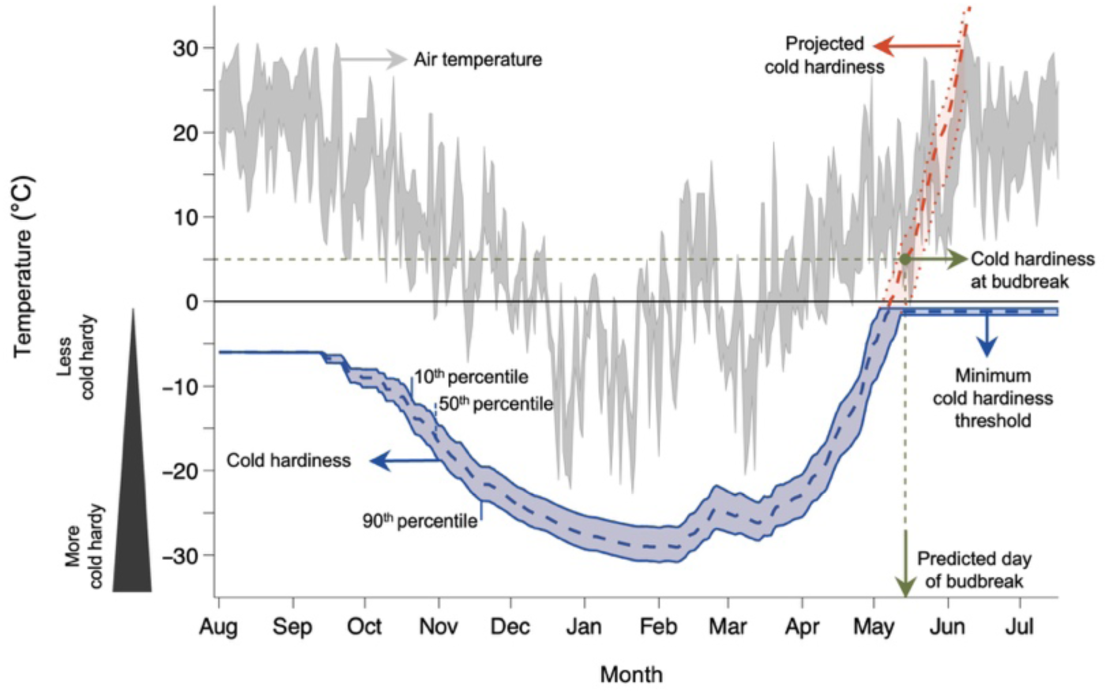
Conceptual overview of the cold hardiness model to predict budbreak. The NYUS.1 cold hardiness model (Kovaleski *et al*., 2023) is used to predict the time to budbreak for *V. labrusca* ‘Concord’ grapes grown in Portland, NY, US in the 1983-1984 dormant season. Cold hardiness (blue ribbon) is gained and lost in response to seasonal fluctuations in air temperature (grey ribbon). Two different end trajectories are shown for the cold hardiness prediction: real cold hardiness values that remain negative in value, where CH stops decreasing once it reaches minima extracted from ^70^ (blue ribbon); and a second, projected cold hardiness where values are calculated into non-freezing temperatures (red ribbon and dashed line). When the projected cold hardiness reaches a threshold determined for cold hardiness at budbreak [established from data in Kovaleski et al. (2018)], budbreak is predicted to occur.

The full phenology dataset contains a total of 329 observations – recorded as the Julian day of year (DOY) of budbreak – spanning a cumulative total of 68 seasons (1955-1956 to 2022-2023). An exhaustive list of locations, cultivars, and observational timespans for each location can be found in **Supporting Information Table S1**. In most cases, phenology data was explicitly reported as the day of year to reach 50% of buds at BBCH 07 and BBCH 09 [Biologische Bundesanstalt, Bundessortenamt und Chemische Industrie; Meier et al. (2009)], or the equivalent stage 4 in the modified Eichhorn-Lorenz scale for grapevines (Coombe, 1995). Occasionally, observations were made of 50% of buds at stage BBCH 05 (and in one location, observations by one observer as first bud at stage BBCH 05).

### Phenology predictions

To predict phenology, we first predicted cold hardiness using the NYUS.1 grapevine cold hardiness model (Kovaleski *et al*., 2023). This model uses temperature data as inputs to generate cold hardiness predictions as outputs.

### Weather Data and Chill Accumulation

Daily maximum and minimum temperature values were retrieved for all eight locations in this study through publicly available databases. For European locations, downscaled (1 km^2^ spatial resolution) data were obtained through the easyclimate package in R (Cruz-Alonso *et al*., 2023) for the period of 1950-2022, which was the most broad timespan available using these resources. In Bordeaux, FR, additional temperature data were accessed from a nearby weather station (station ID: 335500003) for the time period of 1 January 2023 – 30 June 2023.

For two of the three North American locations (Niagara-on-the-Lake, ON, CA; and Lewisburg, PA, US), high resolution (1 km) temperature data were obtained from Daymet version 4 (Thornton *et al*., 2021) and accessed via *daymetr* version 1.7.1 (Hufkens *et al*., 2018) for the period of 1980-2022, the broadest timespan available. For Portland, NY, US, temperature data from the weather station located at the Cornell Lake Erie Research and Extension Laboratory were used (available through the Lake Erie Regional Grape Program) for the time period of 1926-2023. We primarily used gridded climate datasets to ensure consistent, quality-controlled temperature records across sites and years, incorporating station-based observations only where high-resolution, long-term data were available.

For evaluations using general yearly weather, such as sensitivity (see below), we characterized each location:year combination using the mean temperature for the dormant season calculated by obtaining the average temperature from 1 November to 30 April for each season within each location, as this is the period of time when most shifts in cold hardiness occur. This is referred to simply as mean dormant season temperature.

### Modeling Cold Hardiness

The model utilized in this study is the NYUS.1 cold hardiness prediction model (Kovaleski *et al*., 2023) (see **Notes S1** for a detailed description of the model). NYUS.1 integrates (1) *Acclimation* and (2) *Deacclimation* processes over daily time steps from bud set to any given point during the dormant season, using daily temperatures as an input:

1. *Acclimation*: The rate of acclimation for buds at any given day is a function of chilling accumulation, the difference between the daily minimum temperature and the maximum temperature that elicits chilling accumulation on any given day, and the cold hardiness of the previous day. Generally, (i) as chilling accumulates, temperatures must be lower to result in acclimation, (ii) the lower the temperature, the more cold hardiness it can elicit, and (iii) the more cold hardy buds are on the day before, the less cold hardiness they can gain when exposed repeatedly to the same temperature (e.g., a very cold hardy bud exposed to mild temperatures in late winter will not gain any further cold hardiness).
2. *Deacclimation*: The effective rate of deacclimation at any given day is determined by a temperature response, calculated for daily minimum and maximum temperatures, which is modulated by a chilling response curve – the deacclimation potential. Generally, (i) the effective rate of deacclimation is the product of the rate of deacclimation and the deacclimation potential, where (ii) higher temperatures result in higher rates of deacclimation, and (iii) high chilling accumulation leads to higher deacclimation potential (e.g., a bud exposed to high temperatures at low levels of chilling accumulation will not lose any cold hardiness).

Chilling accumulation is an input within NYUS.1. There are many possible models used to calculate chill as a response to temperature [see Wang *et al*. (2020) for 12 examples of models]. The responses to chilling accumulation within NYUS.1 are based on chill portions calculated through the Dynamic Model [Fishman *et al*. (1987a,b); model C_9_ within Wang *et al*. (2020)], which therefore serves as a sub-model. The Dynamic Model was used as there was no reparameterization of the NYUS.1 model, and thus the parameters used to describe dormancy progression are based on chill portions. The Dynamic Model is a complex chilling accumulation model, often left undescribed [e.g., within Wang *et al*. (2020)]. Generally, temperatures between ∼–2 and 13 °C promote chilling accumulation of chill portions. Once a chill portion is accumulated, it is not lost, but warm temperatures can cause loss of a precursor of a chill portion. However, fluctuations between low and warm temperatures can slightly enhance chilling accumulation. Chilling accumulation was calculated in RStudio using *chillR* version 0.75 (Luedeling *et al*., 2023). Since the Dynamic Model calculates chill portions in hourly timesteps, daily minimum and maximum temperatures were transformed into hourly data using the “*stack_hourly_temps*” function in *chillR.* Chilling accumulation for each location was calculated continuously from 1 July to 30 June for every year where temperature data was available. To summarize, the cold hardiness of buds at any given time is calculated as an integration of functions of temperature over time, which are modulated by chilling accumulation, which is itself a response to temperature. In this process, acclimation is the strongest driving force for cold hardiness during the fall, and deacclimation is the strongest driving force during spring, resulting in a general U-shaped curve for cold hardiness.

The 50^th^ percentile cold hardiness predictions used the same parameters originally published in Kovaleski et al. (2023) and are based on field data collected in Geneva, NY, US. To provide additional detail regarding potential damage, we used the same calibration and validation datasets to obtain parameters for the 10^th^ and 90^th^ percentiles for cold hardiness predictions. These parameters are described in **Supporting Information Table S2**.

### Spring Cold Hardiness

Minimum cold hardiness thresholds during spring are not set within the NYUS.1 model, allowing for the projection of cold hardiness freely towards positive temperature values, which here we use for predictions of budbreak (see below). For analyses concerning the estimation of frost damage risk and safety margins, we limited the cold hardiness to thresholds which describe the minimum cold hardiness of growing tissue, obtained from the literature (Gardea, 1987). Such thresholds were obtained by Gardea (1987) through controlled freeze tests, evaluating damage on buds at “fourth flat leaf” stage (BBCH 14) that were exposed to 1°C temperature steps between 0°C and -4°C, and a +4°C control. Here, we assumed that newly emerged tissues in springtime reach a minimal level of cold hardiness, that this level is then maintained constant over time, and that these values for grapevine do not differ on a genotype-specific basis [as suggested by the little effect of continent of origin reported for species within a genus; see Kirchhof et al. (2025)].

### Cold Hardiness at Budbreak

Projected cold hardiness trajectories for each cultivar were used to predict the day of year by which 50% of buds reached the BBCH 07 (“beginning of bud burst: green shoot tips just visible”) stage. Two different thresholds for the cold hardiness at budbreak were applied, based on cultivar: a predicted cold hardiness of +10°C was used for Cabernet-Sauvignon and Riesling, and +5°C was utilized for Concord. Thresholds were determined from data within Kovaleski et al. (2018) in which rates of cold hardiness loss in forced deacclimation assays were related to different bud developmental stages.

Additional threshold adjustments were made to account for differences in observational habits within phenological records. The first adjustment was used to account for years and locations where phenology observations were determined at the 50% BBCH 05 (“wool stage”) stage rather than at BBCH 07. Because buds at BBCH 05 are more cold hardy, we used a +5°C threshold so that the predicted cold hardiness values aligned with the developmental stage represented in the observation for Riesling and Cabernet-Sauvignon (North & Kovaleski, 2024). An additional adjustment was made specifically for predictions in Bordeaux, FR over the 1981-2008 period. During this time, the phenological record for budbreak was the day of year when the first bud for each cultivar reached BBCH 05 (as opposed to 50% of buds at a certain stage as in other locations and years). Because the first bud to reach BBCH 05 corresponds to an earlier developmental stage with substantially lower cold hardiness, we used a threshold of 0°C for Cabernet-Sauvignon and Riesling predictions during the 1981-2008 period to align the model output with the stage represented in the observational habits (see **Supporting Information Fig. S1** for a schematic representation of threshold setting and use).

### Adjustment of Cold Hardiness Predictions to Account for Damage

In years where low temperature damage to buds was predicted, a correction was applied to the cold hardiness of the remaining buds based on an estimate of the degree of damage (as a percentile value). This was achieved through a simple linear interpolation. Damage estimates were performed for days within each season where the minimum air temperature was lower than the predicted cold hardiness. In seasons where multiple instances of damage were predicted, percentile values for damaged buds were calculated for each event, based on the existing population immediately prior. The cold hardiness of the remaining buds was calculated after each damage event, dependent on both the existing cold hardiness percentile values preceding the event and the percentile estimation of buds damaged from the freeze event.

When the minimum air temperature fell between the 10^th^ and 50^th^ percentile cold hardiness values, the following equation was used to estimate the percent of buds damaged from the existing population:

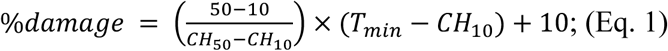

where *T_min_* is the minimum temperature of the day, and *CH_10_* and *CH_50_* are the 10^th^ and 50^th^percentile cold hardiness values. When the minimum air temperature fell between the 50^th^ and 90^th^ percentile cold hardiness values, the following equation was used to estimate the percent of buds damaged from the existing population:

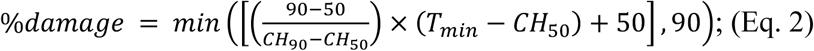

where *CH_50_* and *CH_90_* are the 50^th^ and 90^th^ percentile cold hardiness values, and the constants (e.g. 10, 50 and 90), represent the corresponding quantiles associated with each cold hardiness value. After calculating the estimated percent of buds that were damaged, cold hardiness percentile values were adjusted to account for the buds remaining in the undamaged population using the following equations:

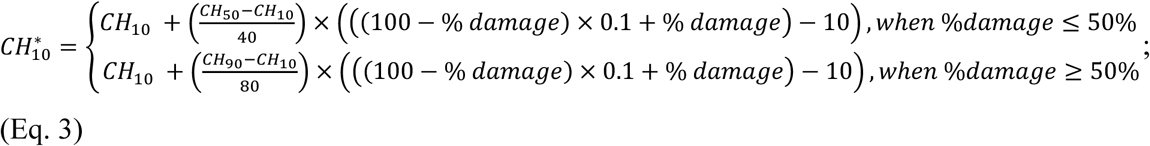

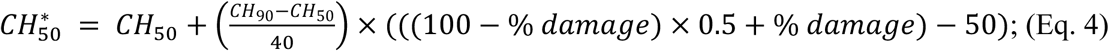

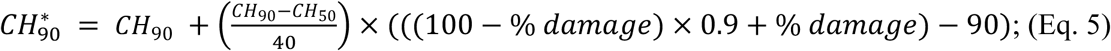

Where 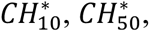 and 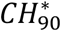 are the damage-adjusted predicted cold hardiness for the 10^th^, 50^th^, and 90^th^ percentiles, respectively. Damage-adjusted cold hardiness predictions were used in all analyses presented unless otherwise mentioned (see **Notes S2** for an example illustrating the application of the equations 1-5 described above).

### Validation of Damage Predictions

Given the absence of long, publicly available records of empirically determined cold hardiness measurements, we validated our predictions of cold damage by comparing them with historical reports. To do so, we performed this analysis based on Concord cold hardiness modeled in one location (Portland, NY) over the entire time period of cold hardiness predictions (1926-2023). Our analysis focused on Concord in the Lake Erie region given its local economic importance, guaranteeing mentions of cold damage in a local news source, and the availability of searchable news reports in English. The historical records used were obtained from two sources: *The Erie Daily Times* (1927-2006) – a local newspaper – and *Veraison to Harvest* (2007-2023) – an extension publication by Cornell University which reports on New York State’s grapevine production. For newspaper articles, the archival search was done using the keywords ‘grape+frost’. In order to be included in the analysis, the articles and extension reports had to meet the following criteria: (i) there was a mention of any degree of frost damage to the grapevine crop or (ii) articles explicitly mentioned that the crop escaped damage. If no records of damage were found for a given year, it was assumed that no damage occurred to the buds.

A positive detection of damage for a season was considered when at any day of the season the daily minimum temperature was lower than the predicted 10^th^ percentile cold hardiness for that day. A binary outcome was then applied to all years in this analysis, regardless of level of damage, or number of damage events, where 1 indicated damage occurrence during a season, and 0 indicated no damage. A binary outcome was also applied to the mention of damage in records, wherein 1 indicated presence (records indicated damage occurred within a season) and 0 indicated absence (records indicated no damage had occurred, or no records of damage were discovered for a given season). These binary outcomes were then used to construct a confusion matrix to evaluate prediction performance and calculate sensitivity, specificity, precision, and accuracy of the model.

### Effect of Chilling Temperature Range

To examine potential errors associated with chilling range on the budbreak predictions, both as a whole and by location, we applied a series of phase shifts to change the relative contribution of temperatures to chilling accumulation in the Dynamic Model. The chilling range was thus shifted by 1°C intervals, from -6°C to +6°C during chill accumulation estimation (e.g., chill is maximized at 7.5 °C within the Dynamic Model, and we shifted this maximum to occur between 1.5 °C and 13.5 °C). Budbreak predictions were then produced using the model without any other adjustments. RMSE for predictions at each phase shift step were then calculated.

### Indicators for Climate Suitability and Sensitivity

To evaluate the temporal nature of freeze risks within a season, we used temperature records from Portland, NY, over the time period of 1926-2023, to predict cold hardiness and subsequently estimate the frequency of freeze events and potential freeze damage. Safety margins defined as the difference between the minimum registered air temperature (*T_min_*) and the 50^th^ percentile cold hardiness prediction were calculated daily (*CH_50_*):

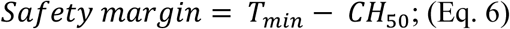

Where a safety margin below zero indicates at least 50% freezing damage is predicted, and positive values indicate different levels of safety. We defined freeze risk as the number of days when the safety margin dropped below zero for each cultivar in all locations for all years in which weather data was available (n=1626 location:cultivar:year combinations). Therefore, safety margin calculations were also used to determine the number of freeze risk injury days over the range of mean dormant season temperatures experienced (November-April), providing an overall measure of how many days during the season there was a risk of damage:

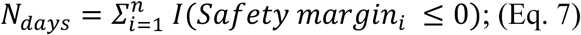

The potential effects of temperature changes on phenological timing (sensitivity) were evaluated by studying the relationship between the predicted day of budbreak and the mean dormant season temperatures over all locations, years and cultivar combinations (n=1626). Differences in responses to different mean dormant season temperatures for the entire dataset, and also in three temperature segments (a) -1°C to 3°C, b) 3.01°C to 10°C, and c) 10.01°C to 12°C) were calculated in RStudio using the *rms* package, version 6.8-1 (Harrell & Frank, 2024).

### Evaluation of budbreak predictions

Accuracy and performance of the budbreak prediction model, and the effect of different variables (i.e., observer, cold damage, and chill accumulation) were evaluated through determination of Bias, Root Mean Square Error (RMSE), and Coefficient of determination (*r*^2^), comparing predicted budbreak day of year to the effectively independent observations of field budbreak day of year. To evaluate bias, we calculated residuals as the difference between predicted and observed budbreak day of year (*Residuals* = *BB_predicted_* – *BB_observed_*). Bias was then defined as the mean of residuals for each group. Here we used predicted minus observed so that negative values indicate early predictions, while positive values indicate late predictions. RMSE was calculated to quantify the overall prediction error, reflecting both the variance and bias of the model.

To evaluate the effect of observational habit changes in Bordeaux, FR, we tested residuals by observer (three observers) using a one-way ANOVA, with Tukey-adjusted post hoc comparisons of means. Residuals were compared by observer before and after applying a new threshold for cold hardiness at budbreak. To evaluate the effect of predicted damage on predictions of budbreak, we compared three groups to each other: “no damage”, “damage uncorrected”, and “damage corrected”. Residuals for each pair were compared using a one-way ANOVA. All statistical analyses were performed using RStudio. A workflow of the analyses performed here is presented in **Supporting Information Fig. S2**.

## Results

### Cold hardiness can be used to predict budbreak phenology across cultivars & locations

We calculated cold hardiness for all locations, and for each year in which we gathered weather data, we used the projected cold hardiness to predict budbreak (**Figs. 1**, **2**). The initial budbreak predictions across locations and cultivars yielded results (n=329, RMSE=7.3 days, bias=-0.8 days, r^2^=0.88; **Fig 3a**) that are comparable to those reported by other phenology modeling studies (García de Cortázar-Atauri *et al*., 2009; Sgubin *et al*., 2018) **(Notes S3).** Given that cold hardiness trajectories were used to determine budbreak timing (**Figs. 2**, **3b**), it was also possible to group predictions based on the presence or absence of freeze damage to buds. Although our initial predictions did not account for effects of cold damage, given the known delay that occurs in budbreak when cold damage occurs in buds (Chamberlain & Wolkovich, 2021; Charrier *et al*., 2021; Muffler *et al*., 2024a) we produced corrected budbreak predictions whenever damage occurred prior to initially predicted budbreak. A freeze damage event was predicted to occur when the minimum air temperature dropped below (i.e., was more negative than) the cold hardiness prediction associated the 10^th^ percentile of cold hardiness distribution on any given day. For each individual freeze event, a simple linear interpolation was used to determine the proportion of buds damaged, and a new 80% confidence interval was calculated to represent the cold hardiness of the remaining population of living buds. Based on the greater cold hardiness in the remaining buds, budbreak predictions shifted to a later date following freeze damage, with the estimated delay dependent on the magnitude of cumulative freeze damage throughout the dormant season (**Fig. 3b**).

**Figure 3.**
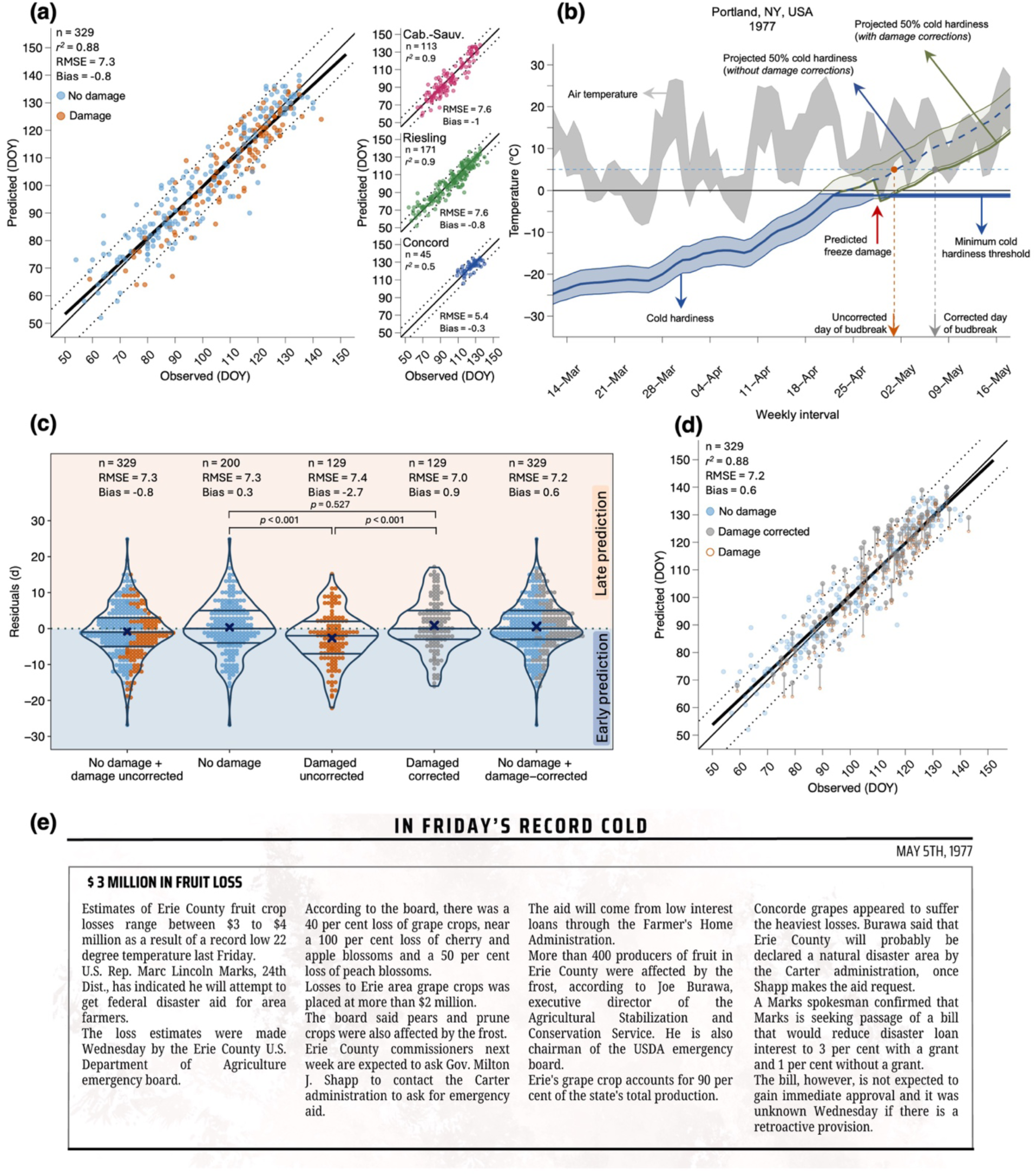
Comparison of budbreak predictions from cold hardiness model with observations. **(a)** Relationship between initial predictions and observations of budbreak. **(b)** Based on expected cold damage prior to budbreak, predictions of budbreak can be corrected, shifting to a later date as the cold hardiness of the surviving bud population is greater. (**c)** Distribution of residuals based on five different groupings, from left to right: initial predictions (years uncorrected for damage + years where damage was not predicted n = 329); years where no damage was predicted (n = 200), uncorrected predictions in years with predicted damage (n = 129), predictions corrected to account for damage (n = 129), and full, corrected predictions (predictions corrected to account for damage + predictions in years where damage is not predicted, n = 329). Residuals for each pair were compared using a one-way ANOVA. (**d)** Comparison of predictions to observations after accounting for the consequences of freeze damage. (**e)** Sample article from historical records used to qualitatively validate damage predictions (Erie Daily Times, 5 May 1977). In (**a,b**), thin solid black line represents 1:1 ratio between observations and predictions, and dotted lines represent an interval of +/- 7 days from 1:1 line. Heavy black line shows linear relationship between predicted and observed points.

Grouping of model residuals based on predictions of freeze damage revealed that errors were significantly influenced by the presence or absence of such damage (No damage vs. Damaged uncorrected; **Fig. 3c**). In years where damage was predicted to occur, uncorrected budbreak predictions were on average 2.7 days earlier than observed budbreak (i.e., Bias_damaged, uncorrected_ =–2.7d), significantly contrasting from years where no damage was predicted (Bias_no damage_=+0.3 days). After accounting for damage in budbreak predictions, error in years where damage was corrected did not significantly differ from years without damage (No damage vs. Damaged corrected; *p* = 0.527). As a result of incorporating a method for predicting and accounting for damage in the budbreak predictions, overall model outputs improved (RMSE=7.2 days, bias=0.6 days, *r*^2^=0.88; **Fig. 3d**).

Because long-term cold hardiness measurements were unavailable, we used historical reports from extension documents and newspapers for one location as a proxy for evaluating the model (**Fig. 3e**). While these records likely underreport damage, they provide the only multi-decadal criterion against which to compare predictions, allowing us to assess whether NYUS.1 captures years with widespread, well-documented events. This approach classified reported damage in the records as a binary variable (damage vs. no damage). Years were classified as either “damage” or “no damage”, where any mention of damage to grapevine buds was determined to be a “damage” year, and years where damage was not mentioned were recorded as “no damage” years. The analysis was performed using Concord cold hardiness predictions in Portland, NY, US over the time period of 1926-2023 (n=97). The binary validation of damage predictions over the time period where cold hardiness predictions were made (1926-2023) resulted in model sensitivity of 0.75, specificity of 0.72, precision of 0.57, and accuracy of 0.73 (**Supporting Information Fig. S3**). Sensitivity and specificity values above 0.70 suggest that the model is reasonably effective at correctly identifying both positive and negative cases, while the lower precision highlights that some false positives remain (Bujang & Adnan, 2016) – although some false positives here may be due to underreporting of damage. Based on the evidence that (i) damage-corrected budbreak predictions better reflect phenological observations and (ii) predictions of damage are also reflected in historical records, corrected predictions were incorporated into all further analyses presented. Hereafter, these are then referred to as freeze damage-corrected budbreak (FDC budbreak).

### Observational habits and chilling as significant sources of error

Using FDC budbreak, we explored the effect of different factors contributing to uncertainty in the model outputs using partial residual plots. We found observer – based on observational habits – to be a significant source of uncertainty (**Supporting Information Fig. S4**). Year did not show a significant effect on residuals, and other factors such as minimum yearly temperature in any given location, maximum predicted cold hardiness, and chill accumulation showed effects in only one cultivar (**Supporting Information Fig. S5**). Importantly, the effect of chilling on residuals showed a positive (albeit not significant for two cultivars) slope for the three cultivars when combined, indicating systematic issues with chill accumulation – be that through modeling of accumulation, or how chilling is incorporated in the cold hardiness model. However, the predictions using NYUS.1 remained robust even under climatic conditions that differ substantially from those where the model was developed – particularly warmer locations with lower chill accumulation (**Fig. 4, Supporting Information Fig. S6**).

**Figure 4.**
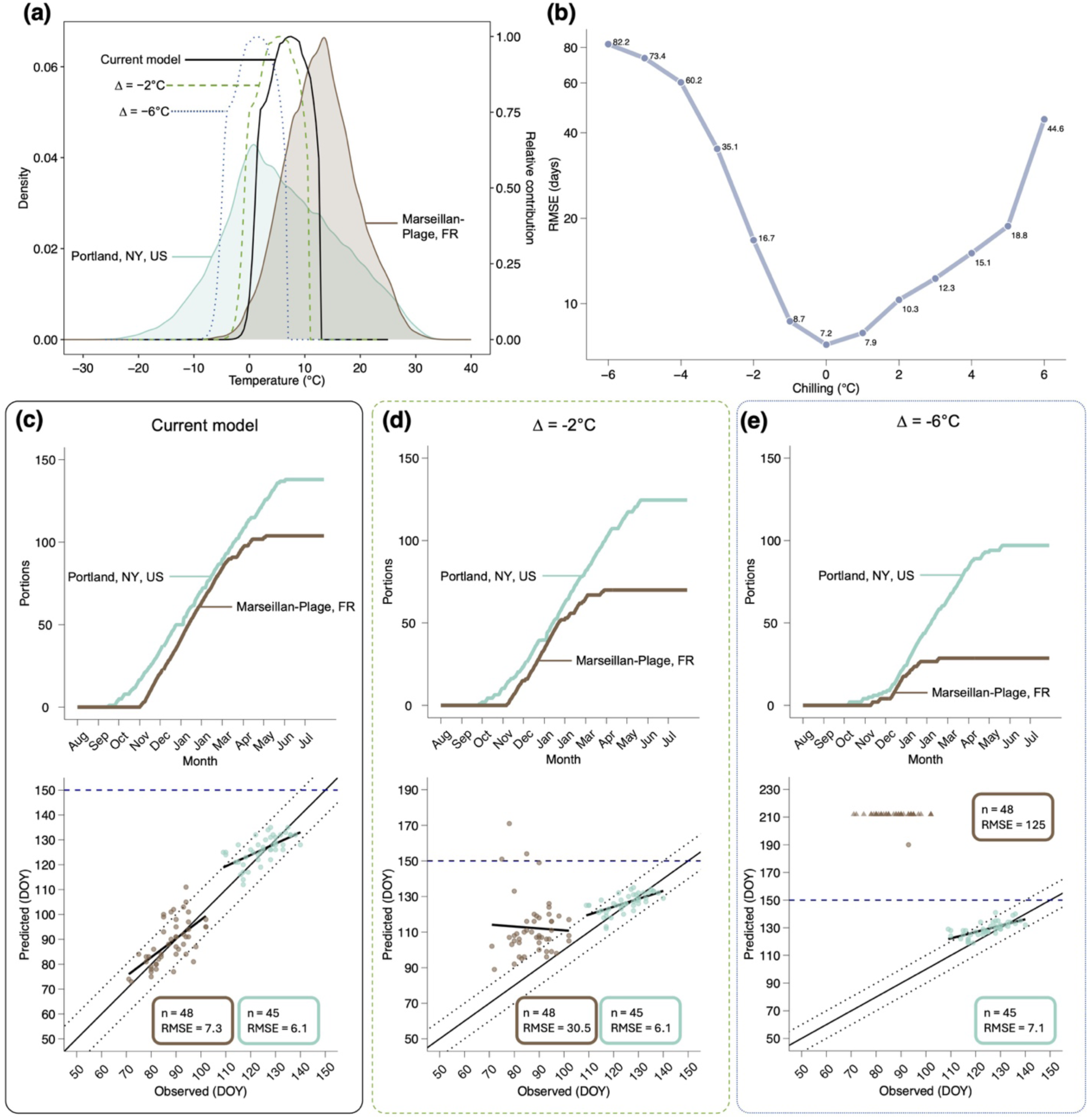
Phenology predictions in different sites are differentially affected by shifts in how chilling accumulation is calculated. (**a)** Density plot of recorded temperatures in Portland, NY, US and Marseillan-Plage, FR, and the relative contribution of temperatures to chilling accumulation in the Dynamic Model under unaltered and two sample phase-shift scenarios (-2°C and -6°C). (**b)** RMSE of budbreak phenology predictions under a range from -6°C to +6°C of phase shift scenarios. **c,d,e,** Phenology predictions compared to observations and average chilling accumulation in a cold location (Portland, NY) and warm location (Marseillan-Plage, FR) under (**c**) unshifted conditions, (**d**) a -2°C phase shift, and (**e**) a -6°C phase shift scenario. In (**c,d,e**), thin solid black line represents 1:1 ratio between observations and predictions, and dotted lines represent an interval of +/- 7 days from 1:1 line. Heavy black line shows linear relationship between predicted and observed points. Horizontal dashed line at DOY 150 in (c,d,e) is used to highlight change in axis in (e), and points shown as an arrowhead at 212 days are failed predictions.

Observed date of budbreak in Bordeaux, FR between 1981-2008 were recorded based on the date of first bud to reach BBCH05, or first bud to break, differing from the rest of the record in which dates were recorded based on 50% budbreak to either BBCH05 or BBCH07. We accounted for these differences by decreasing the cold hardiness threshold at budbreak (see **Supporting Information Fig. S1**), allowing predictions to better reflect the earlier stage of development that better corresponds the observations made, reducing this factor to a non-significant source of error (**Supporting Information Fig. S4**).

Chilling accumulation – calculated via the Dynamic Model (portions) and used to modulate both cold acclimation and deacclimation as responses to dormancy – led to a tendency for late predictions at the lower range found in our datasets (i.e., between 85 and 105 portions, *p*<0.001; **Supporting Information Fig. S5d**). Chill portions are accumulated when temperatures fall within a given range, with the amount of chilling accumulated dependent on the frequency and relative contribution of those temperatures to chilling (**Fig. 4a**). Therefore, chilling accumulation throughout the dormant season varies based on the climate of a region and yearly weather patterns. We explored shifting the range of temperatures contributing to chilling to test whether a simple correction would reduce the model residuals. Both increasing and decreasing the temperature range for chill accumulation increased the RMSE of predictions across all regions (**Fig. 4b**). These changes, however, are uneven when observed within climates. In colder locations, where high levels of chill accumulate within a season due to wide temperature distributions, small shifts in chill accumulation range cause virtually no change in predictions (Portland, NY, US in **Fig. 4c,d**). However, in warmer locations where chill accumulation is lower, small shifts towards colder range of chill accumulation results in significant delays in predicted budbreak (Marseillan-Plage, FR in **Fig. 4c,d**). Large shifts towards colder temperature greatly reduce the amount of chill accumulated in warmer locations, significantly affecting budbreak predictions in warmer locations, whereas these effects remain minor in cold areas (**Fig. 4e**).

### Cold hardiness as a driver of climate suitability

Because cold hardiness is a plastic trait that responds to temperature, shifting temperature regimes will influence cold hardiness dynamics to varying degrees, leaving some genotypes more vulnerable to freeze risk during the winter and early spring months. Therefore, we sought to examine the ways in which a cold hardiness model (here NYUS.1) can be utilized to explore low temperature-related growth risks among the three cultivars. To do so, we expanded cold hardiness and budbreak predictions beyond the year:location:cultivar combinations in which phenology observations were available. Instead, for all locations, we estimated cold hardiness and budbreak for the three cultivars, over the range of time for which weather data was available for each location (1950-2023 for European locations; 1926-2023 for Portland, NY, US; and 1980-2022 for all other North American locations).

We first examined predicted freeze risk amongst the three cultivars over the season for a cold location where the longest range of predictions were available (97 seasons; Portland, NY), and where cold damage is often reported (**Supporting Information Fig. S3**). This was done based on safety margin calculations as the difference between the minimum temperature and the *CH_50_* in any given day (Safety margin = *T_min_* – *CH_50_*). Instances of both predicted damage [Safety margin ≤0)] and near damage (0<Safety margin ≤2) were calculated for each cultivar. All seasons were aligned based on each cultivar’s specific FDC budbreak for a given year, and showed different patterns in freeze risk for each cultivar (**Fig 5a**). Cabernet-Sauvignon faced a higher risk of winter damage (i.e., ∼120 to 80 days before budbreak) and a reduced risk of spring damage (i.e., 10 days prior to and after budbreak) when compared to other cultivars. Conversely, Concord displayed low risk of winter damage, but a greater susceptibility to spring damage compared to the other two cultivars. Riesling showed an intermediate risk for both winter damage and spring damage.

**Figure 5.**
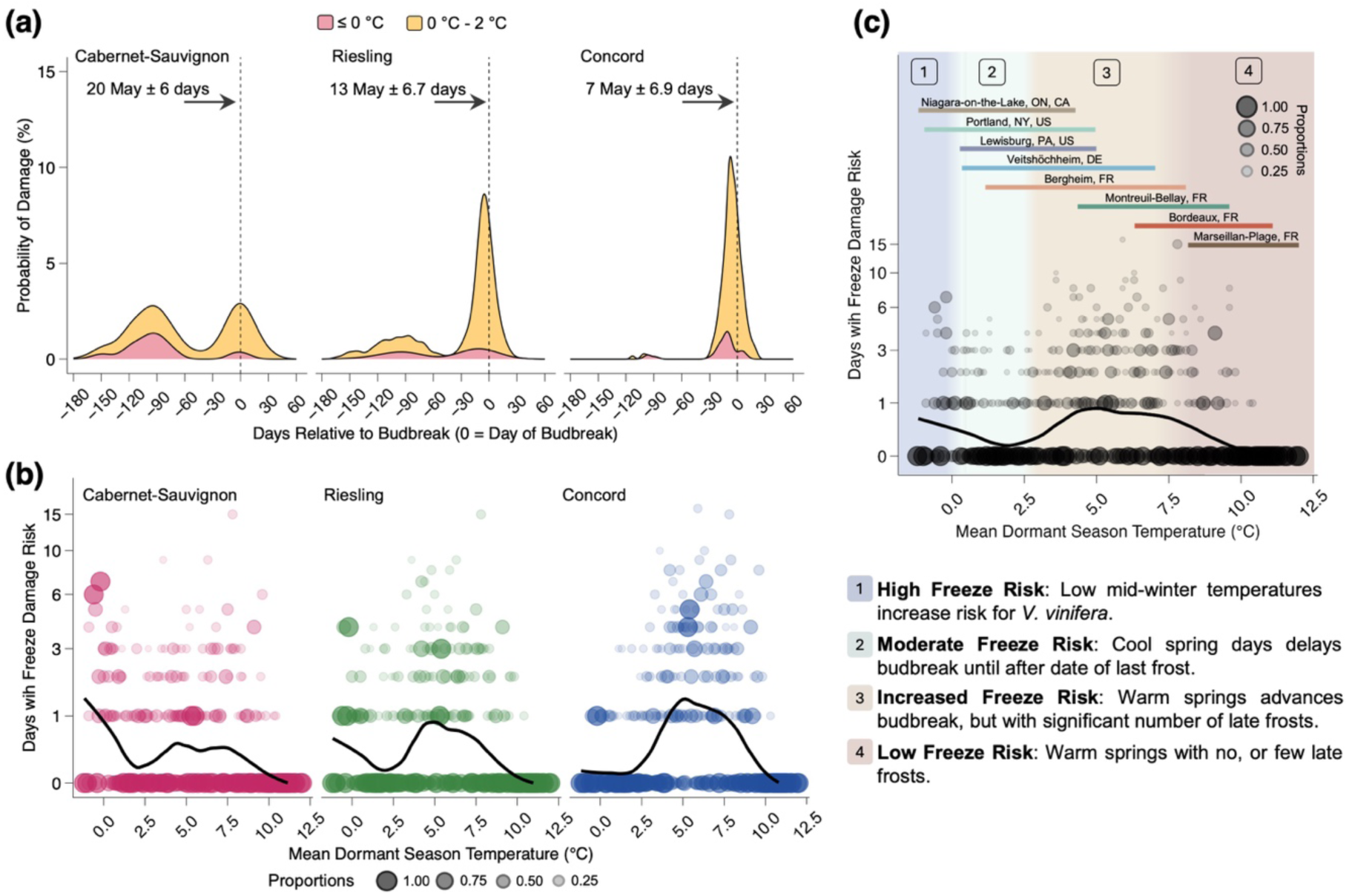
Freeze risk shows temporal, temperature, and cultivar effects. (**a)** Damage risk density plot over time within a season, based on cold hardiness predictions for three grapevine cultivars in Portland, NY, US over the time period of 1927-2023 shows two instances of higher probability of damage: midwinter (120-80 days before budbreak) and around budbreak (15 days before and after budbreak). Red curve represents predicted instances of freeze damage (safety margin less than or equal to 0), and the yellow curve represents instances where there is a narrow safety margin (between 0°C and 2°C). Horizontal axis shows “0” as the mean day of budbreak for each cultivar over the time period. (**b, c)** Number of predicted freeze damage risk days based on mean dormant season temperature (1 November–30 April), for (**b**) each of three grapevine cultivars grown in eight locations and (**c**) for all cultivars combined. Point sizes (**b, c**) are proportional within each decimal mean dormant season temperature. Risk of damage was separated in four temperature bands based on mean dormant season temperature: [1; blue] high freeze risk injury, particularly for *V. vinifera* cultivars, due to low mid-winter temperatures; [2; green] cool spring days delay budbreak until last frost; [3; orange] risk of freeze damage increases as spring phenology advances faster than date of last frost; [4; red] temperatures are too warm for low temperature injury to be significant. Horizontal colored lines (**c**) show mean dormant season temperature ranges in datasets from each of the eight locations studied.

To more easily generalize responses to current and future climates, we explored the influence of dormant season temperature (November-April) on freeze damage risk as the number of days within a season where minimum temperature was lower than the predicted *CH_50_* (negative safety margin). For this, predicted cold hardiness for the three cultivars across the eight locations and for all years was used. Cultivar-dependent patterns in risk became evident: colder dormant seasons (mean temperature below ∼2.5°C) resulted in a greater freeze risk for Cabernet-Sauvignon and (to a slightly lesser degree) Riesling, while Concord exhibited negligible risk (**Fig. 5b**). With moderate temperatures (e.g., 5°C), all cultivars exhibited some level of freeze risk, with risk being highest for Concord, and lower for Riesling, and Cabernet-Sauvignon. When evaluating all cultivars together, we can separate freeze risk broadly into four different temperature bands (**Fig. 5c**). Mean winter and spring temperatures below 1°C were considered high freeze damage risk temperatures. Moderate freeze damage risk was assigned to mean temperatures ranging from 1°C to 3°C. Increased freeze damage risk was identified for temperatures ranging from 3°C to 8°C, and temperatures above 8°C were classified as low freeze damage risk.

We also analyzed the sensitivity of simulated budbreak based on mean temperature of the dormant season using the predicted FDC budbreak for all cultivar:year:location points (**Fig. 6a**). At any mean dormant season temperature, Concord has earlier budbreak, followed by Riesling, and by Cabernet-Sauvignon. Date of budbreak advances with increasing mean dormant season temperatures, with sensitivity of –5.8 days/°C (i.e., 5.8 earlier budbreak DOY with every degree of warming) for all cultivars combined. At the cultivar level, Concord has a slightly faster advancement compared to Riesling and Cabernet-Sauvignon (–6.0 days/°C, –5.5 days/°C, and – 5.7 days/°C, respectively). However, the overall response appears to be non-linear, with varying patterns across different temperature ranges and locations. To highlight these differences, mean winter and spring temperatures in Portland, NY, US (one of the coldest locations in our study) and Marseillan-Plage, FR (the warmest location in our study) were explored (**Fig. 6b**). Portland shows an overall phenology advancement of only –3.3 days/°C, with mean dormant season temperatures ranging from -1°C to 5°C (Fig 6B). This contrasts with Marseillan-Plage, FR, where mean dormant season temperatures range from 8°C to 12°C, showing a shifting pattern within its temperature range. There is a phenology advancement of –6.8 days/°C when mean dormant season temperatures ranges from 8°C to 10°C. However, as mean temperatures increase further in that location, phenology starts to delay at a rate of +3.8 days/°C when mean dormant season temperatures are above 10°C (**Fig. 6b**). When data for all locations is split into three separate groups, sensitivity is highest in moderate temperatures (–6.3 days/°C for 3-10°C), and lowest and positive for warmest temperatures (+2.0 days/°C for 10-12°C; **Fig. 6a**).

**Figure 6.**
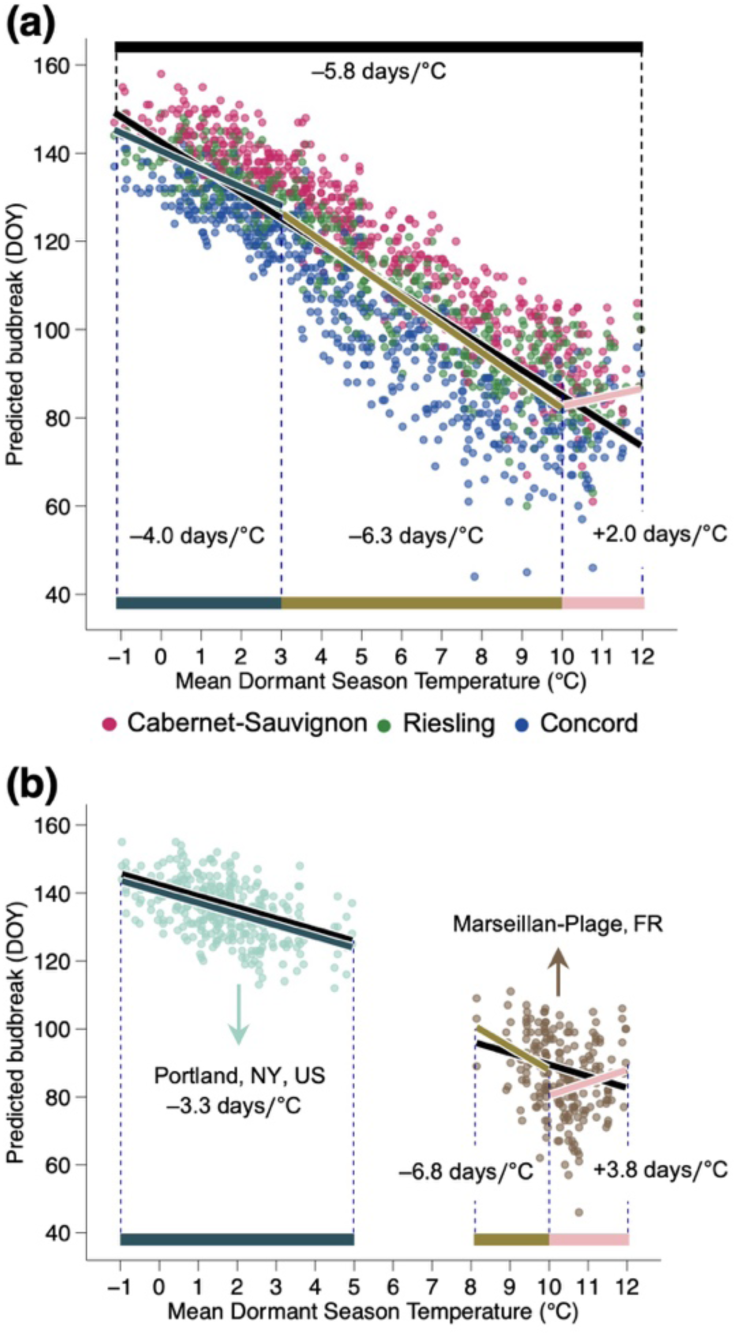
Sensitivity of phenology to temperature is uneven across the range of climates studied. **(a, b)** Freeze damage corrected predicted day of budbreak for all cultivars as a response to mean dormant season temperatures (1 November–30April) for (**a**) all eight locations combined and (**b**) separately for Portland, NY, US and Marseillan-Plage, FR. (**a**) Budbreak advances at an overall rate of 5.8 days/°C for all cultivars, while showing different magnitudes and directions at different temperature intervals, (**b**) which is also affected by local temperature dynamics.

## Discussion

For temperate woody perennials, spring phenology (here budbreak) occurs after plants have been released from an annual period of winter dormancy (Lang *et al*., 1987). Time spent in cold temperatures, quantified as chilling accumulation, is required for dormancy release. Then, to break bud and resume growth, time spent in warm temperatures (forcing) is required (Lang *et al*., 1987; Chuine & Régnière, 2017). Many attempts to model budbreak rely at least partially on this relationship, using quantifiable thermal time metrics to elucidate the time required to reach the stage (Chuine, 2000). Here, the negative chilling and forcing relationship is substituted by winter dynamics of cold acclimation and deacclimation, thus incorporating a third temperature response – cold hardiness – to accurately predict spring phenology while allowing for better understanding of cold temperature-related suitability. Our findings demonstrate that a process-based cold hardiness model for grapevine [NYUS.1; Kovaleski et al. (2023)], trained on cold hardiness data for a single location, is generalizable to predict spring phenology (as budbreak) across a wide range of global climates in which grapevines are typically cultivated (**Figs. 1**,**3**). Our analysis also shows that the NYUS.1 model effectively predicts freeze events (**Supporting Information Fig. S3**), while highlighting genotypic differences in damage patterns across locations and climatic conditions (**Fig. 5**). These results emphasize the importance of incorporating cold hardiness as a predictable trait within a phenology model, offering valuable insights beyond just predicting budbreak that will be consequential as global climate conditions continue to rapidly change.

Cold hardiness has a u-shaped pattern (**Fig. 2**), where its level at any point depends on prior temperatures experienced and specific species or cultivar. Here we use a cold hardiness threshold to predict budbreak. Given that colder climates and colder seasons can elicit greater cold hardiness (**Supporting Information Fig. S7**), different locations can require different degrees of deacclimation (or forcing) to reach budbreak. This variation helps explain why the same genotypes show different chilling and heat requirements when evaluated in different locations. Additionally, reacclimation is possible if colder temperatures occur after a period of deacclimation [i.e., **Fig. 2**, see deacclimation period in late February, followed by reacclimation in early March; Kalberer et al. (2006), Arora & Rowland (2011)]. In this context, reacclimation could be considered analogous to what would be negative forcing, or negative growing degree-days in ordinary spring phenology models – a concept contrary to growth, and thus never used, yet applicable and realistic within a cold hardiness dynamics framework.

The suitability of woody perennial species to local climates is shaped by numerous interacting environmental factors and phenotypic plasticity (Baranger *et al*., 2024). Among these, exposure to low temperatures requires particular attention, especially in light of the unprecedented rise in global temperatures (Masson-Delmotte *et al*., 2021). Increasing temperatures have been shown to cause increased risk of late frosts in some areas (Augspurger, 2013; Sgubin *et al*., 2018; Vitasse *et al*., 2018). Our study found that all cultivars exhibited advancement in phenology with warmer dormant season temperatures (**Fig. 6**), consistent with broader trends in Northern Hemisphere populations (Li et al., 2019; Piao et al., 2019; Fernandez et al., 2023; Li et al., 2024). Here we used mean dormant season temperature (Nov-Apr) as NYUS.1 has a complete overlap of chilling and forcing phases (i.e., acclimation and deacclimation are possible at any time), thus making it not possible to separate temperatures for each phase. However, it is important to note that warmer winter temperatures can also lead to increases in freeze damage potential, and the trends depend on cultivar [**Fig. 5b**; or similarly provenance in Mura et al. (2022)] and where a location lies in terms of mean dormant season temperature (**Fig 5c**). Our study found that a general pattern occurs, where risk generally decreases with increasing mean temperatures, but a “bump” appears between ∼3°C and 8°C. This “bump”, represents a mismatch between phenology advances due to warmer springs and the potential for significant late frosts, increasing the freeze damage risk. When looking at each cultivar separately, the phenology-temperature mismatch places Concord at a greater risk for spring frost damage, including the timing right before budbreak, whereas Riesling and Cabernet-Sauvignon have lower risk (**Fig. 5a,b**). This further demonstrates the advantage of using a cold hardiness model to predict spring phenology, allowing for simultaneous assessments of both phenology advances and freeze damage risk that includes pre-budbreak damage.

In addition to risks related to freeze damage, excessive warming reduces chilling accumulation and delays budbreak based on our predictions (**Figs. 4**,**6**). Recent work supports these findings, suggesting that delays in phenology are also possible due to reduced exposure to temperatures which contribute to chilling accumulation (Kramer *et al*., 2000; Benmoussa *et al*., 2017; Gaeta *et al*., 2018; Li *et al*., 2019; Fernandez *et al*., 2023; Mo *et al*., 2024), something often observed in areas with marginal chilling conditions where temperate woody perennial crops are introduced (Malagi *et al*., 2015; Du *et al*., 2019; Song *et al*., 2020; Zeng *et al*., 2023) and empirically determined in general chilling-forcing experiments [e.g., Flynn & Wolkovich (2018); Kovaleski (2022)]. As changing global temperatures shift spring phenology, woody perennials will face novel risks in their current distributions (Chmura et al., 2011; Ford et al., 2016; Baranger et al., 2024; Stuke et al., 2024). The degree to which phenology shifts among the different genotypes in a given location is ultimately an effect of their plasticity and adaptive capacity on separate aspects of winter physiology: cold hardiness, chilling, and forcing. This highlights the importance of expanding empirical work with cold hardiness and chilling to include diverse genotypes and populations, particularly in warm range edges of natural ecosystems and marginal agricultural conditions, where plants will be more strongly affected (**Fig. 4**).

Grapevine cultivars possess a great diversity in traits that affect responses to climatic conditions (Flynn & Wolkovich, 2018; Morales-Castilla *et al*., 2020). This diversity is evident when exploring cold hardiness dynamics across the cultivars presented in this study. Across all locations, predicted patterns in acclimation and deacclimation reflected known genotypic differences in hardiness and development, and are therefore reflective of current distributions of the genotypes studied (**Fig. 1**). Cabernet-Sauvignon and Riesling – both cultivars of *V. vinifera* – exhibit low and intermediate mid-winter freeze tolerance (**Figs. 5a, S7**), respectively. These cultivars are thus best suited to climates with moderate to mild winters and longer growing seasons that are presently characteristic of many temperate climates with winters moderated by large bodies of water and western European locations. Concord, a cultivar of the more cold hardy *V. labruscana,* showed the greatest freeze tolerance among the three genotypes here, making it best adapted to regions with the cold winters and shorter growing seasons, presently characteristic of many viticultural regions in Northeastern North America. When freeze risk is evaluated based on mean dormant season temperature, Concord shows higher risk at moderate temperatures due to its faster deacclimation and advanced budbreak compared to the other cultivars (**Figs. 5c**, **6a**). Genotypic adaptation to local climates has allowed for successful grapevine cultivation in a wide range of climates, and our assessment of spring damage risk supports these known differences. Understanding the degree to which cold hardiness dynamics respond to fluctuations in dormant season temperatures – and, ultimately, how this influences budbreak phenology – will be imperative to making informed management and planting decisions as local climates continue to change.

Although the framework presented here removes subjectivity in determining the transition between chilling and forcing temperatures, exploring alternative approaches to chill calculation could be beneficial to further improve model outputs. The Dynamic Model has been shown to be one of the best models for comparing chilling across regions (Fernandez *et al*., 2020), and particularly without acknowledging cold hardiness (North & Kovaleski, 2024). NYUS.1 uses chill portions to calculate deacclimation potential as an increasing logistic response that translates into the negative exponential relationship of budbreak time (or forcing) to chilling accumulation (Kovaleski, 2022). Here, small shifts in temperature for chilling accumulation cause large increases in error (**Fig. 4b**). Chilling accumulation is a crucial component of most phenology models, and in NYUS.1, chilling accumulation modulates the magnitude of acclimation and deacclimation responses throughout the season. Given the significance of chilling to the model variability and the expected future declines in chill accumulation in regions like southern France (**Fig. 4c-e**) (Fernandez *et al*., 2023), improving existing chill models or developing new ones is necessary to better reflect physiological responses to cold temperatures (Gaeta *et al*., 2018; North & Kovaleski, 2024). Alternatively, reparametrizing chill models such that they reflect species or genotype-specific behaviors in chill accumulation could also potentially improve the performance of this and other phenology models (North *et al*., 2024). Our work thus adds to a growing body of literature showing that cold hardiness is likely an important factor to be acknowledged in studies of chilling and dormancy, which is further substantiated as even at the molecular level cold hardiness genes have been implicated in mechanisms associated with dormancy [i.e., *CBF* genes directly promoting expression of *DAM/SVL/SVP2*, Ding et al. (2024)].

Although there was no parameter optimization for the cold hardiness model, predictions of budbreak resulted in very good statistics – particularly after correcting for cold damage (RMSE=7.2 days, Bias=0.6 days, *r^2^*=0.88; **Fig. 3**). The NYUS.1 model was developed using only grapevine bud cold hardiness data obtained in Geneva, NY, US (Kovaleski *et al*., 2023), while parameters which relate cold hardiness dynamics to chilling accumulation were obtained from a study of 15 unrelated woody perennial species grown in Boston, MA, US [**Fig. 2** in Kovaleski (2022)]. This demonstrates that process-based models that use physiologically-derived parameters can be used across climates – even if chill accumulation has a small effect on residuals (**Supporting Information Fig. S5d**). Measurements of cold hardiness, particularly in warmer climates, or using cold hardiness-informed chill models could be helpful in removing chill accumulation as a source of error. Our metrics are comparable to or better than studies using different models that included calibration based on budbreak data. Ferguson *et al*. (2014), found a general RMSE of 7.3 days (n=152) for budbreak predictions of 23 grapevine cultivars in Washington state, USA using the WAUS.2 cold hardiness model. However, their predictions had a low coefficient of determination (*r^2^*=0.45), likely due to the little variability in dates of budbreak for a single region. Considering the poor performance for the WAUS.2 model to predict cold hardiness in different locations (Kovaleski *et al*., 2023; Wang *et al*., 2024), we can speculate that budbreak predictions would also be poor. García de Cortázar-Atauri *et al*. (2009) found RMSE ranging from 9.2 to 18.3 days for three models (one with two parameter sets) for Cabernet-Sauvignon (n=63) and Riesling (n=64) in four regions in France. Sgubin *et al*. (2018) found RMSEs between 9.7 and 11.1 days for Cabernet-Sauvignon (n=71) and Riesling (n=65) using two different phenological models in four regions in France. Moreover, we performed a comparison of model performance using PhenoFlex, considered to be one of the strongest established phenology modeling approaches within the chilling-forcing framework [Luedeling *et al*. (2021); see **Supporting Information Figs. S8, S9,** and **Notes S3** for a full explanation]. PhenoFlex was parameterized and tested in two different approaches, all of which returned RMSEs that were comparable to those produced by NYUS.1 (**Notes S3**). The RMSE is assessed by comparing model outputs with observed budbreak dates. Standardized methods for recording phenology observations, such as EL and BBCH scales, are widely used. However, some degree of subjectivity is unavoidable when the phenotyping is based on observations of characteristics that cannot be easily discretely quantified. In multi-decadal phenology records spanning multiple locations with different observers, inconsistencies in observation methods are inevitable (Liu *et al*., 2021). A further limitation is the lack of information on observation frequency, which may constrain the achievable accuracy of model-observation comparisons. Here we demonstrated that, depending on previously known discrepancies in observational habits within phenology records, adjusting the threshold of the NYUS.1 model can improve predictions for budbreak (**Supporting Information Fig. S4**). As has been previously described in other organisms (van Strien *et al*., 2008), our results reinforce that observations using 50% occurrence of different phenological stages have better results than using first observations – even with corrections. With expansion of phenology modeling, the development of standardized, quantifiable methods for observing budbreak, including the use of imaging [e.g., PhenoCam (Katal *et al*., 2022)], would enhance model accuracy across locations, observers, and species – advances that will likely contribute to a deeper understanding of dormant season physiology.

Previous modeling work on climate suitability for grapevines only included a mid-winter threshold (Morales-Castilla *et al*., 2020). However, dynamics of weather are extremely important in midwinter cold hardiness, and therefore some of the potential viticultural expansion into colder climates predicted may be overestimated. Our predictions included a significant amount of years and locations with some level of cold damage (129 out of the 329 total observations). The equations used to estimate damage and correct subsequent cold hardiness trajectories are based on known delays in time to budbreak that occurs as a result of low temperature damage (Evans *et al*., 2019; Chamberlain & Wolkovich, 2021; Muffler *et al*., 2024b). We demonstrated that correcting for predicted cold damage (when temperature drops below cold hardiness levels) can improve predictions of budbreak timing. Importantly, we validated our predictions of damage using historical records in the location with the most extensive temperature record (Portland, NY, US; **Figs. 3, S3**). Besides extensive temperature records for Portland, NY, qualitative records of damage (newspaper records and extension reports) were also longstanding, publicly available, and easily accessible. We also assumed that newspaper records would consistently report damage events, given the economic importance of grapevines in the region. Nonetheless, smaller instances of damage, those occurring pre-budbreak may have gone unreported. This may have resulted in an underrepresentation of damage events, in turn, resulting in a higher false positive rate. Because these historical reports are incomplete and heterogeneous in detail, this component of the validation should be regarded as preliminary, but our analysis indicates these reports may be a possible source of data for cold damage.

Though this analysis does suggest that NYUS.1 can accurately predict damage, further subjectivity exists in the historical records, with variable language used to describe the damage and the degree to which damage was observable. For example, NYUS.1 predicted some instances of only slight damage when records of damage were not found. A likely explanation is that the damage was light and localized, and thus not widely reported, resulting in the more conservative nature of the model based on higher false positive predictions (**Supporting Information Fig. S3**). To further improve these predictions, experiments may be required, including defining minimum cold hardiness thresholds for different tissues (Proebsting *et al*., 1978; Gardea, 1987), and measuring mid-winter cold hardiness in multiple locations – particularly warmer climates where this trait is often easily ignored. It is important to note, however, that cold damage is known to delay spring phenology (Chamberlain & Wolkovich, 2021; Qiu *et al*., 2024; Muffler *et al*., 2024a), and ordinary budbreak models have trained on phenology datasets that contain varying levels of cold damage. Thus, shifts in damage patterns that are expected depending on increasing temperatures [e.g., **Fig. 5** in our study, and Lamichhane (2021)] may be a cause of failure for such models as climate changes, while that is unlikely to affect a model using cold hardiness as its basis.

Here, we present a novel method for predicting spring phenology, producing accurate budbreak predictions using a previously published cold hardiness model (NYUS.1 Kovaleski *et al*., 2023), without calibrating its parameters to the budbreak dataset used in this study. Such phenological predictions can be used for planning of agricultural practices related to different phenological stages, such as sprays for pest control and pruning. Furthermore, we also fit new NYUS.1 parameters for different damage percentiles, allowing predicted cold damage (as a result of temperatures dropping below modeled cold hardiness) to shift budbreak predictions and thereby improve their accuracy.

The good predictions without calibration indicate that the physiologically derived parameters used within the NYUS.1 framework work well in non-analogous climates, despite having originated from data for a single location. Through instances where reacclimation occurs, we use a physiological basis to introduce the equivalent of negative forcing into phenological modeling. Beyond budbreak, our framework for predictions includes cold hardiness, allowing for corrections of predictions based on damage incurred, as well as assessments of cultivar suitability to different locations and climates, both presently and in future conditions based on winter physiology. Improvements of winter physiological responses are still possible, such as better defining chilling accumulation effects. Therefore, empirical determination of dormancy and cold hardiness in different climatic gradients are warranted. Continued work in measuring cold hardiness of many species and in varied climates [e.g., Charrier et al. (2013); Deslauriers et al. (2021); Kovaleski (2022)] for empirical validation of cold hardiness predictions will allow for expansion of cold hardiness and budbreak modeling using a framework such as the one presented here, particularly in warmer, marginal climates for temperate woody perennials. Future work may then integrate cold hardiness dynamics into large-scale predictive models to improve forecasts of plant responses to climate change and niche modeling. These efforts can ultimately enable more reliable predictions in earth system models, and better understanding of climatic feedbacks in shifting spring phenology of woody perennials.

## Acknowledgements

We thank M. Chen for comments on the work. This work is partially supported by the National Institute of Food and Agriculture, United States Department of Agriculture, McIntire Stennis projects 7002619 and 1027327, and by the Office of the Vice Chancellor for Research at the University of Wisconsin-Madison with funding from the Wisconsin Alumni Research Foundation.

## Data availability

The data and code for analyses and figures are available through GitHub (https://github.com/apkovaleski/VitisPhenology)

## Supporting Information

**Table S1.** Overview of phenological records used in this study.

**Table S2.** Parameters used for the NYUS.1 cold hardiness model.

**Figure S1.** Conceptual framework of grapevine bud cold hardiness dynamics, deacclimation, and phenological thresholds.

**Figure S2.** Workflow illustration of analysis of cold hardiness and phenology predictions.

**Figure S3.** Confusion matrix for phenology predictions.

**Figure S4.** Effects of observational habits on phenology predictions.

**Figure S5.** Partial residual plots for possible sources of variability in model outputs.

**Figure S6.** Residuals of budbreak predictions by location.

**Figure S7.** Maximum cold hardiness achieved in a season decreases with warmer temperatures.

**Figure S8.** Predicted budbreak using PhenoFlex, trained on a single location.

**Figure S9.** Predicted budbreak using PhenoFlex, trained on the entire dataset.

**Notes S1.** The NYUS.1 model used to estimate cold hardiness for each cultivar.

**Notes S2.** Adjustment of cold hardiness predictions to account for damage.

**Notes S3.** Comparison of NYUS.1 model outputs with PhenoFlex.

