## Supporting Information for "Cold hardiness dynamics predict budbreak and associated low temperature threats in grapevine"

### Contents

**Table S1.** Overview of phenological records used in this study.
**Table S2.** Parameters used for the NYUS.1 cold hardiness model.
**Figure S1.** Conceptual framework of grapevine bud cold hardiness dynamics, deacclimation, and phenological thresholds.
**Figure S2.** Workflow illustration of analysis of cold hardiness and phenology predictions.
**Figure S3.** Confusion matrix for phenology predictions.
**Figure S4.** Effects of observational habits on phenology predictions.
**Figure S5.** Partial residual plots for possible sources of variability in model outputs.
**Figure S6.** Residuals of budbreak predictions by location.
**Figure S7.** Maximum cold hardiness achieved in a season decreases with warmer temperatures.
**Figure S8.** Predicted budbreak using PhenoFlex, trained on a single location.
**Figure S9.** Predicted budbreak using PhenoFlex, trained on the entire dataset.
**Notes S1.** The NYUS.1 model used to estimate cold hardiness for each cultivar.
**Notes S2.** Adjustment of Cold Hardiness Predictions to Account for Damage.
**Notes S3.** Comparison of NYUS.1 model outputs with PhenoFlex.

**Table S1. Overview of phenological records used in this study.** In total, records for the cultivars of interest (*Vitis vinifera* cvs. Cabernet-Sauvignon and Riesling, and *Vitis labruscana* cv. Concord) were obtained for eight different locations and variable climates. Among the records found in the locations used, there is a cumulative total of 329 phenological observations spanning 68 seasons (1955-1956 to 2022-2023).

| Location | Location (in code) | Cultivar(s) | BBCH | Latitude | Longitude | Records Present | Source | Köppen-Geiger Classification |
|---|---|---|---|---|---|---|---|---|
| INRA Colmar Domaine de Bergheim, Bergheim, Haut-Rhin, Grand Est (France) | Colmar | Cabernet-Sauvignon, Riesling | 07 | 48.2053 | 7.3643 | Cabernet-Sauvignon: 1958-1971; 1976-1990<br>Riesling: 1958-2022 | Maury et al. 2023 | Cfb |
| INRA Bordeaux - Villenave d'Ornon, Gironde, Nouvelle-Aquitaine (France) | Bordeaux | Cabernet-Sauvignon, Riesling | 05, 07 | 44.7857 | -0.5762 | Cabernet-Sauvignon: 1970-1982; 1984-1986; 1988; 1995-2008; 2012-2023<br>Riesling: 1970-2008; 2012-2023 | Maury et al. 2023 | Cfb |
| INRA Domaine de Vassal, Marseillan-Plage, Occitanie (France) | Vassal | Cabernet-Sauvignon, Riesling | 05, 07 | 43.3375* | 3.5707* | Cabernet-Sauvignon: 1956-1962; 1964-1967; 1972-1974; 1976; 1988-1997; 1999-2007<br>Riesling: 1956-1965; 1974;1976; 1989; 1996 | Maury et al. 2023 | Csa |
| INRA Domaine de Montreuil-Bellay, Montreuil-Bellay, Maine-et-Loire, Pays de la Loire (France) | MB | Cabernet-Sauvignon, Riesling | 07 | 47.1334 | -0.1377 | Cabernet-Sauvignon: 1982; 1984-1990<br>Riesling: 1982-1986; 1990 | Maury et al. 2023 | Cfb |
| Landesanstalt für Weinbau und Gartenbau (LWG), Veitshöchheim (Germany) | Lower_Franconia | Riesling | 09 | 49.8000 | 9.933 | 1968-1973; 1975-1979; 1981-1984; 1986-1994; 2000-2003; 2005; 2007-2008; 2010 | Bock et al. 2011 | Cfb |
| Portland, NY (USA) | Portland_NY | Concord | 07 | 42.3736 | -79.486 | 1979-2023 | | Dfb |
| Unspecified commercial vineyard in Lewisburg, PA (USA) | Lewisburg_PA | Riesling | 5 (EL), ~07 BBCH | 40.9500 | -76.8833 | 2018-2019 | Persico et al. 2021 | Dfb |
| Four Mile Creek VQA sub-appelation, Niagara Penninsula, Ontario (Canada) | Ontario | Riesling | 4 (EL), ~07 BBCH | 43.2020 | -79.1479 | 2016-2018 | Hérbert-Haché et al. 2021 | Dfb |

*Due to limitations in gridded weather data in proximity to water bodies, coordinates used to obtain weather data for Domaine de Vassal were 43.3375, 3.5485.

**Table S2. Parameters used for the NYUS.1 cold hardiness model.** Parameter combinations used to calculate the 10th, 50th, and 90th percentile cold hardiness predictions for the three cultivars of interest. Parameters for the 50th percentile predictions were based on field data collected in Geneva, NY, US as detailed in Kovaleski (2023). To provide additional detail regarding potential damage, we used the same calibration and validation datasets to obtain parameters for the 10th and 90th percentiles for cold hardiness predictions, which we used to estimate a 80% confidence interval for the predictions.

| Cultivar | $T_{th,low}$ | $T_{th,high}$ | b | c | $CH_{abs\ max}$ | $CH_{bud\ set}$ | d | f | g | h | $k_{deacc}$ | a |
|---|---|---|---|---|---|---|---|---|---|---|---|---|
| Concord [10%] | 0 | 10 | 11 | 66 | -28 | -6 | 3 | 4 | -2.3 | 24 | 1.9 | 3.5 |
| Concord [50%] | 0 | 12 | 8 | 66 | -31 | -6 | 3 | 5 | -2.1 | 24 | 1.9 | 3.5 |
| Concord [90%] | 0 | 12 | 8 | 66 | -33 | -6 | 2 | 4 | -2.2 | 24 | 1.9 | 4 |
| Cabernet-Sauvignon [10%] | 0 | 6 | 6 | 66 | -22 | -6 | 7 | 3 | -1.7 | 14 | 1.7 | 4 |
| Cabernet-Sauvignon [50%] | 0 | 8 | 7 | 66 | -23 | -6 | 6 | 3 | -2.2 | 22 | 1.8 | 5 |
| Cabernet-Sauvignon [90%] | 0 | 10 | 6 | 66 | -25 | -6 | 4 | 3 | -1.9 | 24 | 1.7 | 5 |
| Riesling [10%] | 0 | 10 | 6 | 66 | -24 | -6 | 3 | 5 | -2 | 14 | 2.2 | 5 |
| Riesling [50%] | 0 | 12 | 7 | 66 | -26 | -6 | 3 | 4 | -1.9 | 24 | 2.2 | 5 |
| Riesling [90%] | 0 | 14 | 10 | 66 | -29 | -6 | 2 | 5 | -1.9 | 24 | 2 | 5 |

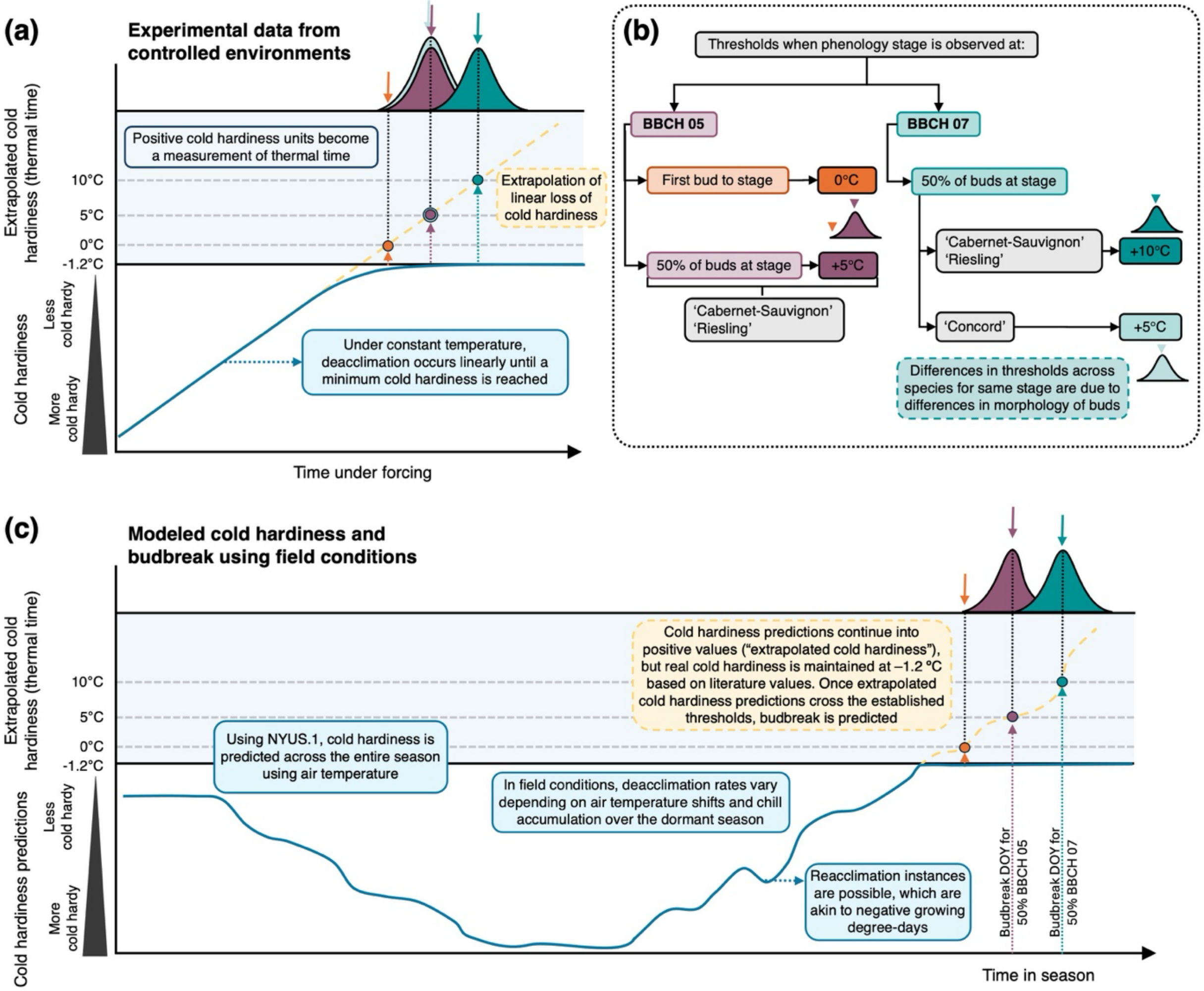


**Figure S1. Conceptual framework of grapevine bud cold hardiness dynamics, deacclimation, and phenological thresholds. (a)** Controlled-environment studies show that under constant temperature, loss of cold hardiness in grapevine buds proceeds linearly until a minimum cold hardiness level is reached (Kovaleski *et al.*, 2018; Kovaleski *et al.*, 2019; North *et al.*, 2022; Londo & Kovaleski, 2025); beyond this point, positive cold hardiness units represent thermal time (extrapolated cold hardiness loss), while tissues largely maintain a level of cold hardiness (Gardea, 1987). Based on the intersection of time to reach a certain phenological stage, and linear model for cold hardiness, a cold hardiness at certain stage can be estimated [see Box 1 within North & Kovaleski (2024) for further information]. **(b)** Thresholds for phenological stages BBCH 05 and BBCH 07 differ among cultivars due to variation in bud morphology (Kovaleski *et al.*, 2019) and were defined for the first bud reaching BBCH 05 or 50% of buds reaching BBCH 05 or BBCH 07 based on data from Kovaleski *et al.* (2018). **(c)** Modeled cold hardiness under field conditions illustrates how estimated cold hardiness values were used to predict budbreak for comparison with historical phenological records.

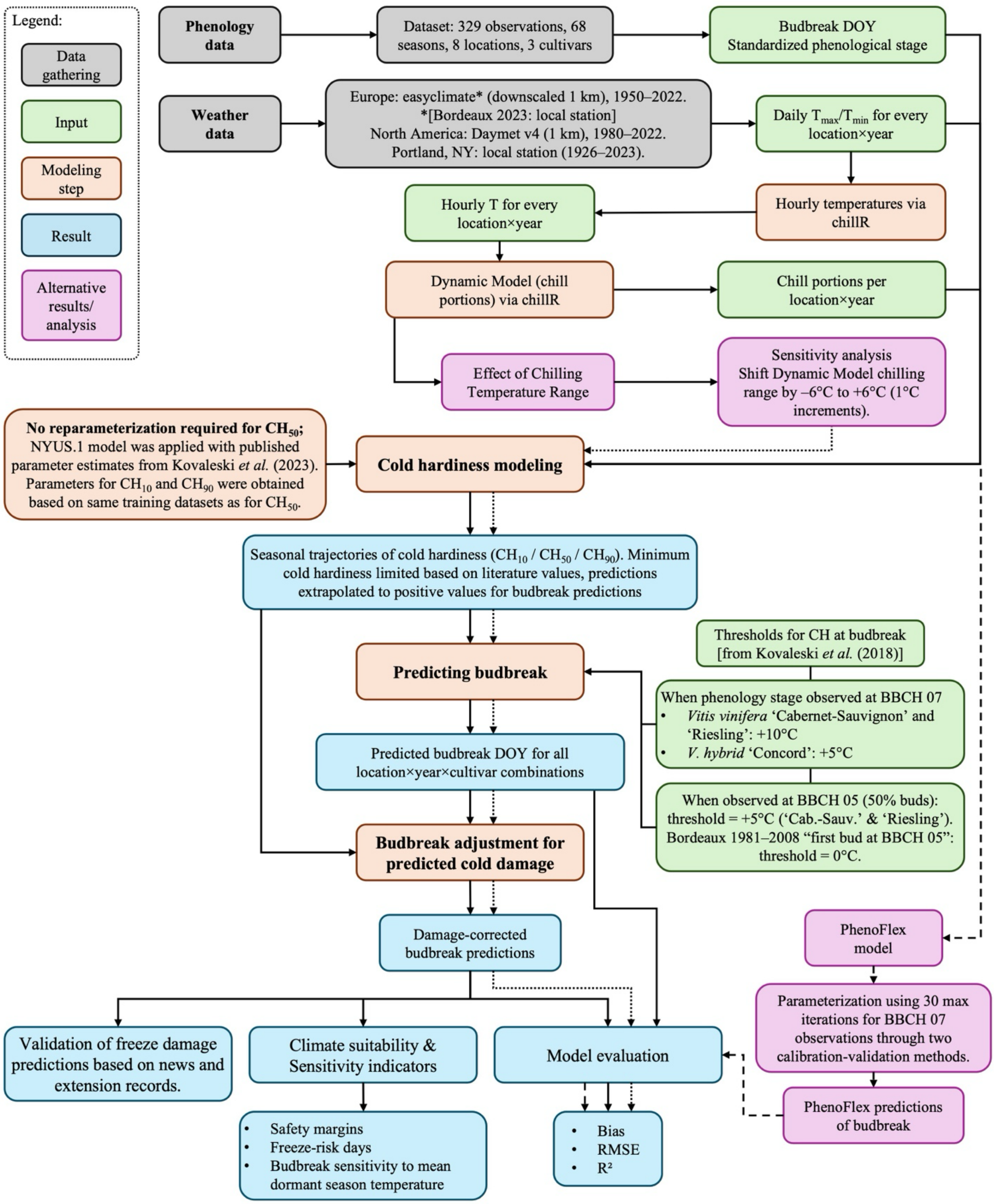


**Figure S2. Workflow illustration of analysis of cold hardiness and phenology predictions.**

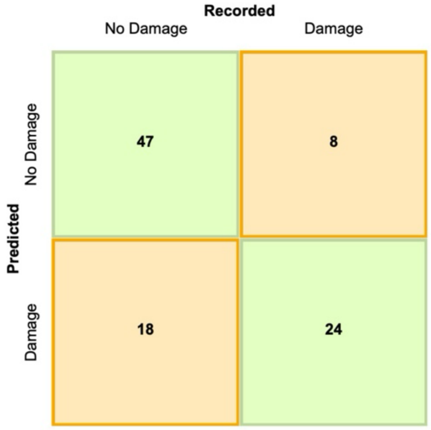


**Figure S3. Confusion matrix for phenology predictions.** Confusion matrix for NYUS.1 model in predicting Concord bud damage compared to mentions of damage in historical records, 1926-2023 (n = 97). Binary outcomes were applied to both the predictions of damage and occurrences in the historical records, where 1 indicated the presence of damage in a season, regardless of the severity of the damage or the frequency of events, and 0 indicated an absence of damage. Sensitivity = 0.75, specificity = 0.72, precision = 0.57, accuracy = 0.73.

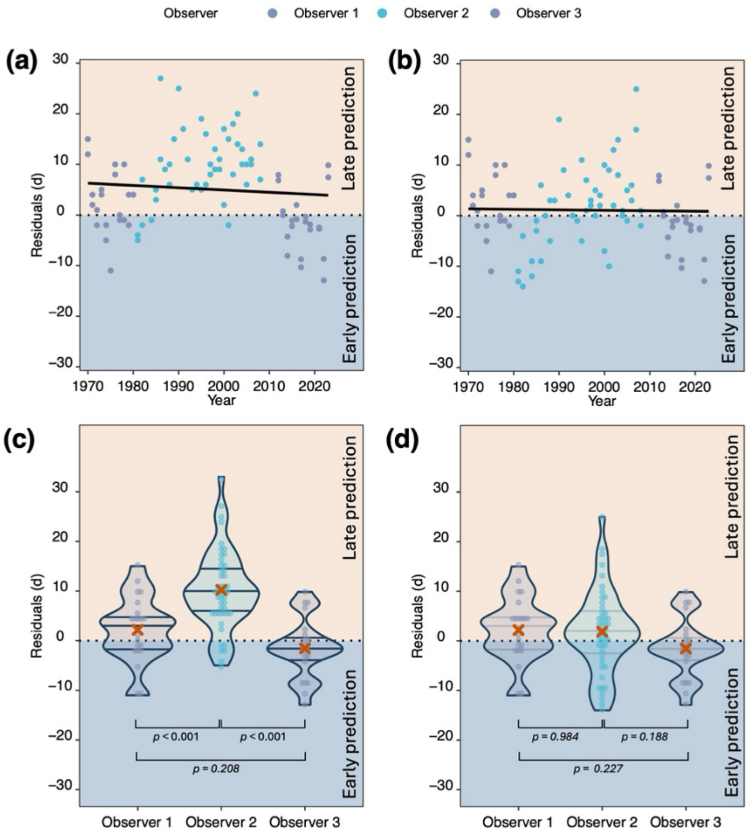


**Figure S4. Effects of observational habits on phenology predictions.** Within the 1970-2019 record of budbreak phenology for Bordeaux, FR, observer 2 used a different protocol compared to observers 1 and 3: observer 2 (1981-2008) defined day of budbreak at first bud to reach BBCH 05; observers 1 and 3 (1970-1980 and 2009-2019) defined day of budbreak when 50% of buds reached BBCH05. Residuals across time show the systematic differences between observational habits when **(a)** these differences are not considered in predictions and **(b)** when these differences are acknowledged within the projected cold hardiness at budbreak coefficient. When comparing residuals between observers, **(c)** observer 2 has significantly different residuals compared to observers 1 and 3 – as NYUS.1 predictions are for 50% budbreak, the predictions are late, **(d)** but the differences disappear when a new projected cold hardiness at budbreak coefficient is used, taking into account the "first bud to reach BBCH 05" observations. **(c,d)** *p* values obtained from Tukey-adjusted post hoc comparisons of estimated marginal means.

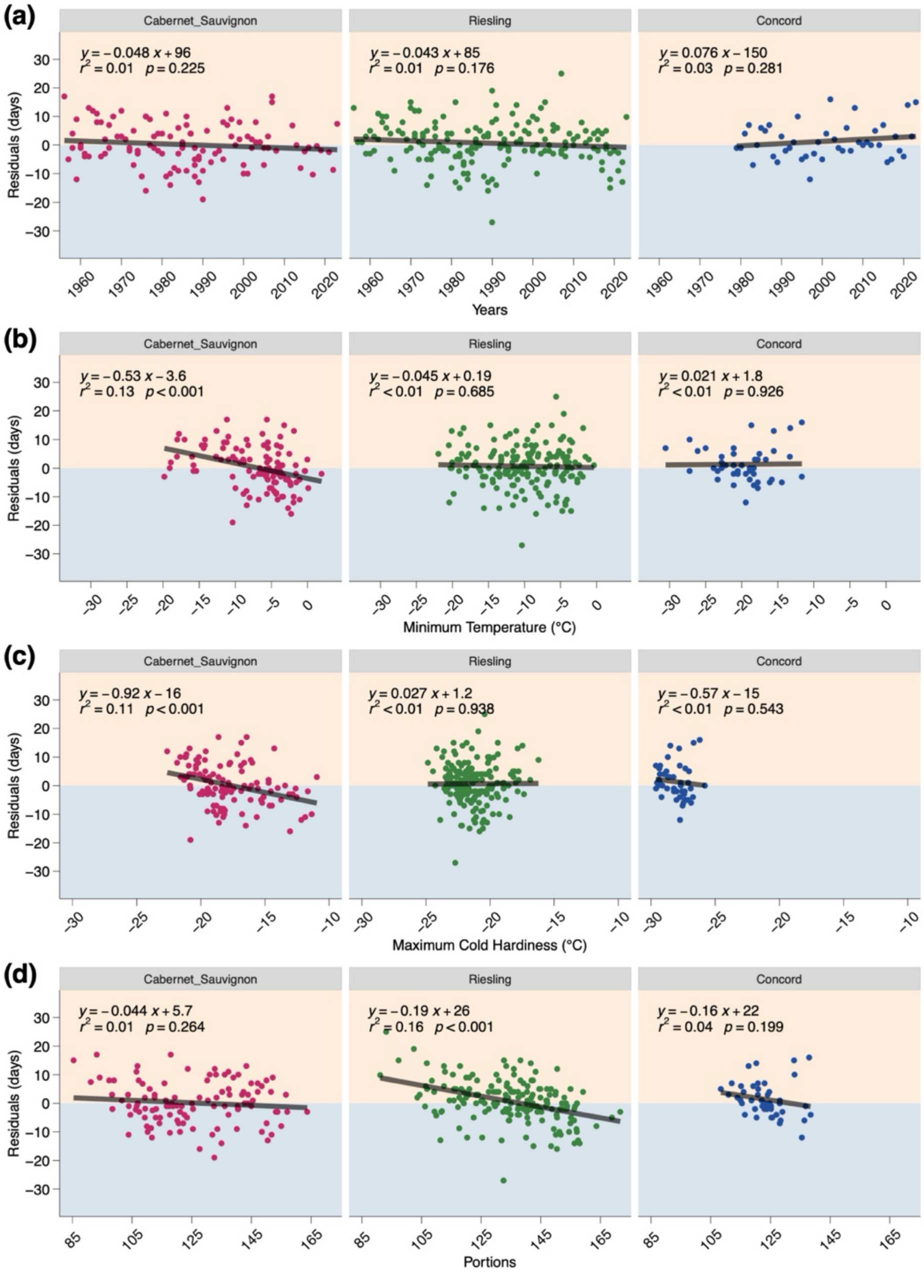


**Figure S5. Partial residual plots by cultivar for possible sources of variability in model outputs.** Using freeze damage-corrected (FDC) budbreak predictions, partial residual plots were produced based on **(a)** time, **(b)** minimum temperature observed in a given season, **(c)** maximum predicted cold hardiness, **(d)** chill accumulation. Fitted lines were generated using simple linear regression.

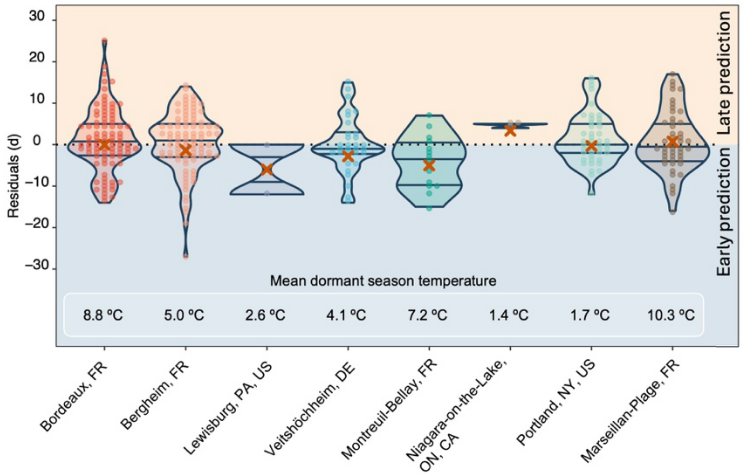


**Figure S6. Residuals of budbreak predictions by location.** Violin plots show the distribution of residuals (observed-predicted) for budbreak day of year across sites, using the FDC predictions. Average mean dormant season temperature (Nov-Apr; °C) for each location in which budbreak phenology was predicted is shown for each location. Orange crosses indicate mean residuals, and black lines show the median and interquartile range.

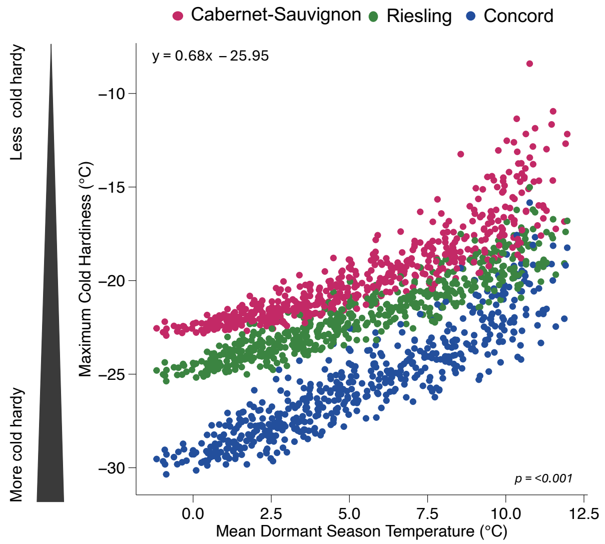


**Figure S7. Maximum cold hardiness achieved in a season decreases with warmer temperatures.** Regardless of genotype, colder climates elicit greater cold hardiness, meaning the same cultivar grown in different locations may require different degrees of deacclimation to reach budbreak—accounting for differences in chilling and heat requirements that are shown for the same genotype in different locations.

**Notes S1.** The NYUS.1 model used to estimate cold hardiness for each cultivar.

### Modeling Cold Hardiness

The model utilized in this study is based on the cold hardiness (*CH*, in °C; **Eq. S1**) prediction model described by Kovaleski et al. (2023) and the generalized *CH* function proposed by Kovaleski (2022) where the cold hardiness integrates (1) *Acclimation* and (2) *Deacclimation* processes that occur over daily time steps from bud set to any given time during the dormant season using temperature records as an input. Kovaleski et al. (2023) describes the process used in this study in detail, including illustrated examples of each equation, however, we present the functions used in brief here. All parameter estimates used are available in **Table S2**.

Overall, the *CH* at any time $t_i$ during the dormant season can be described as:

$$CH_{t_i} = CH_{bud\ set} + \int_{bud\ set}^{t_i} Acclimation\ dt + \int_{bud\ set}^{t_i} Deacclimation\ dt \qquad \text{(Eq. S1)}$$

where $CH_{bud\ set}$ is the initial cold hardiness of buds upon setting in late summer and fall [fixed at –6ºC, the limit of detection that is possible through Differential Thermal Analysis (DTA)], and *Acclimation* and *Deacclimation* are comprised of the functions described briefly below:

(1) *Acclimation:* Acclimation refers to parts of cold hardiness change that are related to gains in cold hardiness. Within NYUS.1, three responses are used to calculate acclimation: one to chill, one to temperature, and on to time. The first is the dependency on chill accumulation, which gives a temperature threshold eliciting gain in cold hardiness, the temperature threshold for acclimation ($T_{th_{acc}}$, in °C; **Eq. S2**). Based on the difference between the minimum temperature in any given day to $T_{th_{acc}}$ of the same day ($\Delta T_{acc}$, in °C; **Eq. S3**), the maximum possible cold hardiness that buds can obtain from that temperature stimulus can be calculated ($CH_{max}$, in °C; **Eq. S4**). To reach a certain $CH_{max}$, exposure to $\Delta T_{acc}$ for a certain amount of time is required. Because the temperature varies constantly in field conditions, this is calculated as an artificial cold hardiness ($CH^*$, in °C; **Eq. S5**) as a response to artificial time ($t_a$, in days) under exposure to the same $\Delta T_{acc}$. Solving **Eq. S5** for the cold hardiness of the previous day ($CH_{t_{i-1}}$), the artificial time for that level of cold hardiness is found. The derivative of **Eq. S5** gives the daily rate of acclimation ($k_{acc}$ in °C day$^{-1}$; **Eq. S6**). Using the artificial time from solving **Eq. S5** as an input in **Eq. S6** provides the rate of acclimation for day *i*. Because the cold hardiness changes are calculated as daily steps, the rate of acclimation is the total gain in cold hardiness for that day. Each of these functions is described in more detail below. The temperature threshold below which acclimation is expected to occur ($T_{th_{acc}}$) is a response to chilling accumulation, represented by a log-logistic curve:

$$T_{th_{acc}}(chill) = T_{th_{low}} + \frac{(T_{th_{high}} - T_{th_{low}})}{1+e^{[b(\ln(chill)-\ln(c)]}} \qquad \text{(Eq. S2)}$$

where the high ($T_{th_{high}}$, in °C) and low ($T_{th_{low}}$, in °C) temperature thresholds, the slope (*b*) and inflection point (*c*) are estimated parameters, while *chill* is the amount of chilling accumulated. NYUS.1 uses the Dynamic Model for chill calculation. Based on Eq. S2, as chilling accumulates, lower temperatures are required for acclimation. The decrease from high to low threshold that is represented here reflects expected physiological changes that occur during chill accumulation and dormancy progression that prioritize deacclimation over acclimation as winter progresses. Note that the slope (*b*) and inflection point (*c*) are shared between this function and the deacclimation potential function ($\Psi_{deacc}$, see "(2) *Deacclimation*" below).

The response to temperature in acclimation is the maximum cold hardiness possible ($CH_{max}$, in °C) elicited by the difference between the minimum temperature on a given day and the acclimation temperature threshold ($\Delta T_{acc}$) as a log-logistic curve:

$$\Delta T_{acc_i}(chill, T_{min}) = T_{th_{acc_i}} - T_{min_i} \quad \text{(Eq. S3)}$$

where $\Delta T_{acc}$ on day *i* is the difference between the temperature threshold for acclimation and the minimum temperature of day *i*. $\Delta T_{acc}$ is a function of chill through $T_{th_{acc}}$. $\Delta T_{acc}$ is an input to calculate the maximum cold hardiness that can result from temperature exposure ($CH_{max}$):

$$CH_{max}(\Delta T_{acc}\,(chill, T_{min})) = CH_{bud\ set} + \frac{(CH_{abs\ max} - CH_{bud\ set})}{1+e^{[d\,(\ln(\Delta T_{acc})-\ln(f))]}} \quad \text{(Eq. S4)}$$

$CH_{bud\ set}$ is the minimum cold hardiness in °C of buds upon maturation and bud set (described previously). The maximum cold hardiness observed over any season is characterized by $CH_{abs\ max}$ and varies by cultivar. *d* and *f* are parameters of the log-logistic curve, wherein *d* represents the slope of the curve, and *f* describes the inflection point.

The next functions describe the rate of acclimation – a response to time – based on the difference between the cold hardiness achieved on the previous day, and the maximum cold hardiness that could be elicited given $\Delta T_{acc}$. For simplicity, this function is separated into two steps. The first step involves calculating the amount of time it would take for buds to reach $CH_{max}$ from $CH_{bud\ set}$, if buds are exposed to constant temperatures. Because exposure to constant temperatures is not reflective of field conditions, time and cold hardiness in this function are classified as artificial. For this an artificial cold hardiness $CH^*$ is calculated as a function of artificial time under exposure to a certain level of $\Delta T_{acc}$:

$$CH^*(t_a, \Delta T_{acc}\,(chill, T_{min})) = CH_{bud\ set} + \frac{(CH_{max} - CH_{bud\ set})}{1+e^{\{g\,[\ln(t_a)-\ln(h)]\}}} \quad \text{(Eq. S5)}$$

where $t_a$ *is the artificial time g* and *h* are the slope and inflection point associated to the log-logistic curve, respectively. $CH^*$ is a function of $\Delta T_{acc}$ through the effect of $\Delta T_{acc}$ on $CH_{max}$. Once the amount of artificial time required to reach $CH_{max}$ has been determined, the rate of acclimation ($k_{acc}$) can be calculated for any time in the field. The rate of acclimation is determined by taking the derivative of the log-logistic function described previously, relating $k_{acc}$ to artificial time.

$$k_{acc}(t_a, \Delta T_{acc}\,(chill, T_{min})) = (CH_{max} - CH_{bud\,set}) \times \left\{0.001 + \frac{\left(e^{\{g\,[\ln(t_a)-\ln(h)]\}}\right)\times\left(\frac{g}{t_a}\right)}{\left(1+e^{\{g\,[\ln(t_a)-\ln(h)]\}}\right)^2}\right\}$$

(Eq. S6)

In brief, the rate of acclimation for buds in the field at any given day is a function of chilling accumulation, the difference between the daily minimum temperature and the maximum temperature that elicits chilling accumulation on that given day, and the cold hardiness of the previous day.

*(2) Deacclimation:* The deacclimation rate is determined by a temperature response curve based on functions from empirical experiments developed by Kovaleski et al., (2023) and Kovaleski (2022) using minimum and maximum daily temperatures as inputs.

For a given day, the rate of deacclimation is a function of temperatures experienced ($k_{deacc}$, in °C day$^{-1}$; **Eq. S7**), calculated using a function adapted from O'Neil *et al.* (1972) for an enzymatic response to temperature for photosynthesis:

$$k_{deacc}(T) = k_{deacc_{max}} \left(\frac{T_{high}-T}{T_{high}-T_{opt}}\right)^a \times e^{\left[a\left\{1-\left(\frac{T_{high}-T}{T_{high}-T_{opt}}\right)\right\}\right]} \quad \text{(Eq. S7)}$$

where $k_{deacc_{max}}$ (in °C day$^{-1}$)is the maximum rate of deacclimation at the optimal temperature $T_{opt}$ (in °C); $a$ is the coefficient related to the slope of the curve, $T_{opt}$; and $T_{high}$ is the temperature at which deacclimation rate becomes zero. $T_{high}$ and $T_{opt}$ were fixed at 40 ºC and 25 ºC respectively.

The deacclimation potential ($\Psi_{deacc}$, unitless proportion, varying from 0 to 1; **Eq. S8**) modulates the amount of deacclimation that actually occurs from the maximum possible, depending on the amount of chilling accumulated as described by a log-logistic response:

$$\Psi_{deacc}(chill) = \frac{1}{1+e^{[-b\,\{\ln(chill)-\ln(c)\}]}} \quad \text{(Eq. S8)}$$

Where $b$ and $c$ are the slope and inflection point of the log-logistic curve, and chill is the chilling accumulation as Dynamic Portions for NYUS.1. The effective rate of deacclimation at any time $t_i$ is then determined as the product of the average deacclimation rate for maximum and minimum daily temperatures ($T_{min}$ and $T_{max}$, respectively) and the deacclimation potential:

$$k^*_{deacc_{t_i}}(T_{min}, T_{max}, chill) = \frac{[k_{deacc}(T_{min}) + k_{deacc}(T_{max})]}{2} \times \Psi_{deacc} \quad \text{(Eq. S9)}$$

In brief, both acclimation and deacclimation are quantified here as responses to daily temperatures and chilling accumulation, which in itself is a response to temperature.

Nine parameters were optimized for each cultivar and are described in Kovaleski et al. (2023), in which a stepwise iterative method was used to minimize the root mean square error (RMSE) between predicted and observed cold hardiness for a calibration dataset produced in Geneva, NY, US. This study uses these previously optimized parameters–which are characterized for 50th percentile cold hardiness predictions–to calculate cold hardiness across all locations presented. For this study, this process was repeated to optimize parameters representing the 10th and 90th percentile cold hardiness estimations. These parameters are described in **Table S2.**

**Notes S2.** Adjustment of Cold Hardiness Predictions to Account for Damage.

Using a simple linear interpolation, we adjusted cold hardiness predictions on days when buds experience low temperature damage (when the minimum air temperature dropped below the predicted cold hardiness threshold). Damage estimates were calculated for each day in the season when damage was predicted.

Based on the pre-damage population, the "new" cold hardiness of the surviving buds was recalculated after each damage event. For example, consider the following case using the equations described in the main manuscript (**Eq 1-5**). Assume the 10th, 50th, and 90th percentile cold hardiness values are –15 °C, –20 °C, and –25 °C, respectively. If the minimum daily temperature on a given winter day falls to –20 °C, we can assume that there was some damage to the buds. To account for this and estimate the "new" cold hardiness of the remaining population, equations (1 and 2) are applied to calculate the percentage of damage and then equations 3, 4, and 5 were used to recalculate the distribution of the 10th, 50th, and 90th percentiles.

Calculating the $\%damage$:

$$\%damage \; = \left(\frac{50-10}{CH_{50}-CH_{10}}\right) \times (T_{min} - CH_{10}) + 10$$

$$\%damage \; = \left(\frac{50-10}{-20-(-15)}\right) \times (-20-(-15)) + 10$$

$$\%damage \; = -8 \times -5 + 10$$

$$\%damage \; = 50$$

In this case, either equation 1 or 2 will yield the same result, since the damage corresponds to 50% of the bud population. After estimating the proportion of buds damaged, the cold hardiness percentile values were then adjusted to reflect the distribution of the surviving (undamaged) population using the following equations. Here we assume that the same distribution continues to exist in cold hardiness (rather than a skewed distribution when a percentage of the buds are damaged).

Calculating the new 10$^{th}$ percentile:

$$CH_{10}^{*} = \begin{cases} CH_{10} + \left(\frac{CH_{50}-CH_{10}}{40}\right) \times \Big(\big((100-\%\,damage) \times 0.1 + \%\,damage\big) - 10\Big), when\ \%damage \leq 50\% \\ CH_{10} + \left(\frac{CH_{90}-CH_{10}}{80}\right) \times \Big(\big((100-\%\,damage) \times 0.1 + \%\,damage\big) - 10\Big), when\ \%damage \geq 50\% \end{cases}$$

$$CH_{10}^{*} \; = -15 + \left(\tfrac{-20-(-15)}{40}\right) \times (((100-10\,) \times 0.5 + 10) - 10)$$

$$CH_{10}^{*} \; = -15 + -0.125 \times (55 - 10)$$

$$CH_{10}^{*} \; = -15 - 0.125 \times 45$$

$$CH_{10}^{*} \; = -15 - 5.625$$

$$CH_{10}^{*} \; = -20.625$$

Calculating the new 50th percentile:

$$CH_{50}^{*} = CH_{50} + \left(\frac{CH_{90} - CH_{50}}{40}\right) \times (((100 - \%\,damage) \times 0.5 + \%\,damage) - 50)$$

$$CH_{50}^{*} = -20 + \left(\frac{-25-(-20)}{40}\right) \times (((100 - 50\,) \times 0.5 + 50) - 50)$$

$$CH_{50}^{*} = -20 + -0.125 \times (75 - 50)$$

$$CH_{50}^{*} = -20 - 0.125 \times 25$$

$$CH_{50}^{*} = -20 - 3.125$$

$$CH_{50}^{*} = -23.125$$

Calculating the new 90th percentile:

$$CH_{90}^{*} = CH_{90} + \left(\frac{CH_{90} - CH_{50}}{40}\right) \times (((100 - \%\,damage) \times 0.9 + \%\,damage) - 90)$$

$$CH_{90}^{*} = -25 + \left(\frac{-25-(-20)}{40}\right) \times (((100 - 90\,) \times 0.5 + 90) - 90)$$

$$CH_{90}^{*} = -25 + -0.125 \times (95 - 90)$$

$$CH_{90}^{*} = -25 - 0.125 \times 25$$

$$CH_{90}^{*} = -25 - 0.625$$

$$CH_{90}^{*} = -25.625$$

Where $CH_{10}^{*}$, $CH_{50}^{*}$, and $CH_{90}^{*}$ are the damage-adjusted predicted cold hardiness (“new” cold hardiness) for the 10th, 50th, and 90th percentiles, respectively.

**Notes S3.** Comparison of NYUS.1 model outputs with PhenoFlex

To provide greater context to the capacity of NYUS.1 to predict budbreak, we compared model outputs to those produced by a classical chilling-forcing approach, PhenoFlex (Luedeling et al., 2021). PhenoFlex is a phenology modeling framework that has been shown to outperform many other existing chilling-forcing models (Luedeling et al., 2021; Tang et al., 2024). In brief, PhenoFlex is distinct from other models because of its ability to characterize the transition from chilling accumulation (calculated with the Dynamic Model) to heat accumulation (calculated with a Growing Degree Hour model) using a sigmoidal transition function. Here, we fit PhenoFlex to the budbreak dataset, following the optimization process approximately as described by Fernandez et al. (2022), through two different calibration-validation methods and compared outputs with NYUS.1. In both methods, we used 30 maximum iterations within the phenologyFitter() function within chillR. To produce the best fitting, we also used a for-loop with set.seed() between 1 and 30, and report the values here where the RMSE of validation were optimized.

In the first calibration-validation method, we fit PhenoFlex using a similar approach as that which occurred for predicting cold hardiness and budbreak in NYUS.1. NYUS.1 was parameterized (i.e., trained) in a cold-winter location (Geneva, NY) for each of the three cultivars (Kovaleski et al., 2023), and these parameters were then applied to model cold hardiness dynamics throughout the season across all locations in the present study without additional training (the entire budbreak dataset is a validation dataset). To simulate this approach, we fit PhenoFlex in a cold location for each cultivar in which data were available, and then applied these parameters across the remaining locations. In the location used for fitting PhenoFlex, the model was parameterized on the entire dataset for the given cultivar, only for observations made at budbreak (BBCH 07 and BBCH 09). Because only one location contained Concord budbreak data, we reserved Concord data only for the next method. PhenoFlex parameters for Riesling and Cabernet-Sauvignon were fit in a cool-climate location in northeastern France (Bergheim, FR), and produced strong budbreak predictions within this location, with RMSE values of 6.2d for Riesling (n = 65) and 5.8d for Cabernet-Sauvignon (n = 29). When these parameters were applied to the remaining locations, prediction accuracy marginally decreased, producing an RMSE for the validation data of 7.3d (n = 117), with cultivar-specific values of 7.0d for Riesling (n = 69), and 7.8d for Cabernet-Sauvignon (n = 48) (**Supporting Information Fig. S8**).

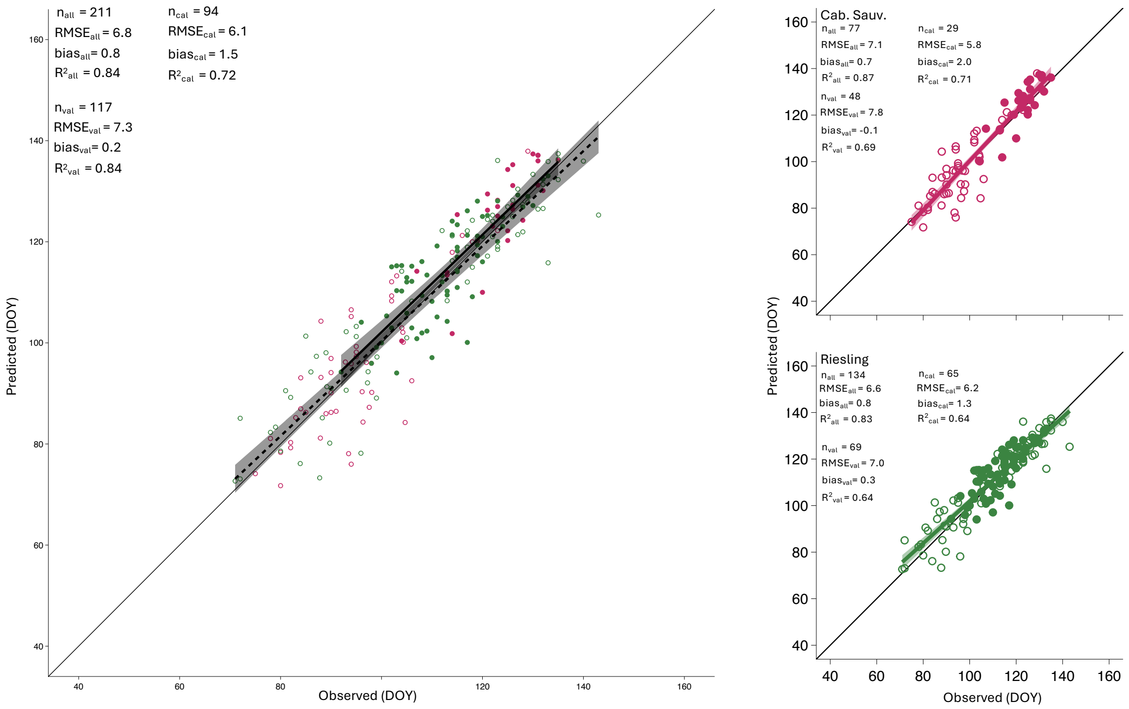


**Figure S8. Predicted budbreak using PhenoFlex, trained on a single location.** PhenoFlex was fitted using the entire dataset for each cultivar in a cool-climate location [Bergheim, FR, for Cabernet-Sauvignon (magenta) and Riesling (green)]), and parameters were then applied to predict budbreak across all remaining locations. $RMSE_{all}$ & $bias_{all}$ = RMSE and bias of predictions from combined fitting and validation datasets. $RMSE_{val}$ & $bias_{val}$ = RMSE and bias of predictions in validation dataset only. $RMSE_{cal}$ & $RMSE_{cal}$ = RMSE and bias of predictions in calibration dataset only. Closed circles represent points used in model calibration, open circles represent points used in model validation.

In the second method, we fit PhenoFlex to the entire budbreak dataset available for each cultivar, again only for observations of BBCH 07 and BBCH 09. This process involved fitting PhenoFlex to a location-naïve phenology dataset to produce a general set of parameters for each cultivar that could be applied across all location:year combinations, using a process similar to the one described in Picornell et al. (2025), though without specifying starting parameters. Using the entire budbreak dataset to train the model gives PhenoFlex the best possible, lowest RMSE. When predictions were compared to phenology observations, PhenoFlex produced an overall RMSE of 6.1d (n = 256), with cultivar-specific RMSE values of 4.6d (Concord, n = 45), 6.4d (Riesling, n = 134), and 6.4d (Cabernet-Sauvignon, n = 77) (**Supporting Information Fig. S9**).

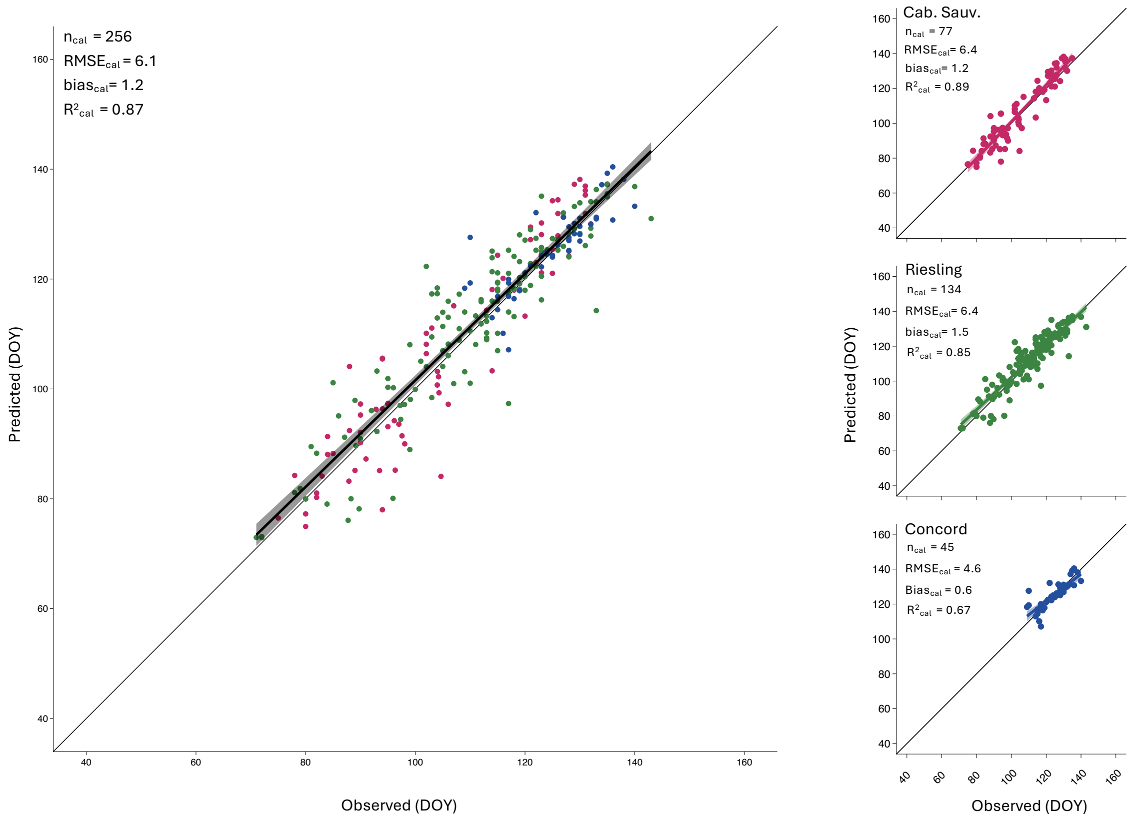


**Figure S9. Predicted budbreak using PhenoFlex, trained on the entire dataset.** PhenoFlex was calibrated using the entire phenology dataset, for all cultivars across all locations. Because there is no separate validation dataset, the reported $RMSE_{cal}$ and $bias_{cal}$ represent the RMSE and bias for the entire dataset and each cultivar, respectively.

For only the observations of BBCH 07 & BBCH 09 (i.e., same dataset used for PhenoFlex), NYUS.1 returns comparable RMSEs, with an overall value of 7.0d, and cultivar-specific values of 6.1d (Concord), 7.4d (Cabernet-Sauvignon), and 6.9d (Riesling). Importantly, however, NYUS.1 produced these predictions *without* being fit to the locations in this study or the same type of data – NYUS.1 was trained on cold hardiness data, but not spring phenology. Essentially, NYUS.1 remains entirely blind to the conditions of each location, and is still able to produce phenology predictions that are on par with those produced even when PhenoFlex is allowed to optimize parameters to fit the entire dataset for each cultivar.
